# Localized network hyperconnectivity leads to hyperexcitability after injury

**DOI:** 10.1101/2022.08.14.503888

**Authors:** Shabnam Ghiasvand, Zubayer Ibne Ferdous, Monika Buczak, Md Fayad Hasan, Yevgeny Berdichevsky

## Abstract

Brain injury increases the risk of the development of epilepsy. Axonal sprouting and synaptogenesis are homeostatically mediated responses by neurons to injury-induced de-afferentation and de-efferentation. Rewiring which occurs due to axonal sprouting and synaptogenesis may alter network excitability and lead to epilepsy. Excitatory and inhibitory connectivity are both subject to homeostatic rewiring. Thus, post-sprouting hyperexcitability cannot be simply explained as result of altered inhibitory/excitatory balance. In this work, we show computationally and experimentally that hyperconnected local networks are created by homeostatic rewiring near the injury site. These local networks are characterized by altered system dynamics despite preservation of excitatory/inhibitory balance. Hyperconnected local networks have a lower threshold for burst initiation, and generate spontaneous bursts, which in turn ignite seizure-like activity in the larger network. Our findings demonstrate a novel network mechanism of hyperexcitability and seizure generation due to maladaptive recovery after injury.

## Introduction

Brain injury increases the risk of the development of epilepsy^1, 2^. Severe brain injury, such as penetrating head trauma, is associated with particularly high risk of epilepsy^3^. One of the mechanisms that may link injury and epilepsy is neural network reorganization, which at cellular level is represented by axon sprouting and synaptogenesis^4^. Axon sprouting has been observed in the hippocampus of patients with temporal lobe epilepsy, and in animal models of acquired chronic epilepsy^5–9^.

Axon sprouting can be induced by injury that involves axotomy and deafferentation^10–13^. The brain reorganizes in response to a wide array of pathologies in order to recover normal function^14–16^. Sprouting has also been observed in the peripheral nervous system. Maladaptive reorganization of peripheral axons after injury may lead to involuntary muscle contractions^17^ or neuropathic pain^18^. Axon sprouting and synaptogenesis in cortex and hippocampus can also be maladaptive. Reorganization of the mossy fiber pathway increased recurrent connectivity in the hippocampus of epileptic patients^5, 6^. Transection of axons of pyramidal neurons in CA3 subregion of the hippocampus resulted in axon sprouting and hyperexcitability^19, 20^. Partial isolation of a region of neocortex led to reorganization of excitatory connectivity of layer V neurons, and emergence of spontaneous epileptiform bursts^21^. Sprouting of excitatory axons may cause hyperexcitability by creating or increasing excitatory feedback. However, sprouting mossy fibers innervate not only excitatory granule cells, but also inhibitory parvalbumin-expressing interneurons^22, 23^. Excitatory synaptic drive to hilar inhibitory neurons increased after injury^24^. Surviving somatostatin inhibitory interneurons also sprout axons and form new synapses between hilus and granule cells^25^, and between CA1 and granule cells^26^. Therefore, inhibitory feedback rises in parallel with increases in excitatory feedback after injury, suggesting that hyperexcitability is not simply a consequence of increased excitatory drive to surviving neurons.

To conceptually understand the development of hyperexcitability, we can consider the case of a pyramidal neuron located close to the site of injury (such as stroke or penetrative head trauma) that results in formation of significant scar tissue or a liquid filled cavity. Pyramidal neurons in intact tissue receive tens of thousands of excitatory and thousands of inhibitory inputs^27^. Most of excitatory inputs are long distance, as the probability of finding an excitatory synapse between two neighboring pyramidal cells is low. After a nearby injury, surviving pyramidal neuron will lose a substantial percentage of its connections, and will therefore have a strong homeostatic drive to re-establish connectivity^28^. However, axons sprouted by this neuron will not find a permissive extracellular milieu with guidance clues that facilitate long-range pathfinding during development. Instead, axons may form synapses with nearby cells which are also experiencing a similar homeostatic drive. Multiple synaptic contacts between pairs of neurons may be made^29^, increasing the strength of the functional connection between them. The resulting connectivity may preserve the balance of excitatory and inhibitory drive to individual cells, when drive is defined as a sum of excitatory or inhibitory synaptic (connection) weights^30^. However, the resulting network will be characterized by hyperconnectivity between cells in a small neighborhood near the injury site (perilesional area). Most of the excitatory input to these cells will be from their immediate neighbors. The dynamics of this local network may make it more hyperexcitable and lead to epilepsy. Here, we investigate this hypothesis by using experimental and computational approaches.

## Results

### Synaptic scaling and hyperexcitability in confined cortical cultures

Dissociated cortical neurons sprout axons and form networks linked with functional synapses in vitro. We created networks of dissociated cortical cells in the presence or in the absence of geometric confinement (Fig. 1(A, B)). We then recorded evoked excitatory and inhibitory postsynaptic potentials (EPSPs and IPSPs, respectively, Fig. 1(C)). There were significantly more connections between pairs of neurons in confined network (60% of *n* = 80 pairs) than in a control network (35% of *n* = 62 pairs) (Fig. 1(H)). Both EPSP and IPSP had larger magnitudes in confined cultures (*n* = 42 and 6 cells, respectively) compared to control cultures (*n* = 19 and 3 cells, respectively) (Fig. 1(D-G)). We also found that stimulation of a single action potential in one neuron in the confined network was more likely to invoke a network burst than single neuron stimulation in control network (Fig. 1(I, J)). At least one burst was evoked in 41% pairs in confined network, compared to 10% of pairs in the control network (*n* = 98 and 62 pairs, respectively) (Fig. 1(K)). Reliability of burst evocation was also higher in the confined network (Fig. 1(L)). These results showed that geometric confinement of sprouting axons (by an artificial physical barrier) resulted in increased probability of connections between neurons, and in scaling up of the strength of both excitatory and inhibitory synapses. These changes in network organization were correlated with hyperexcitability, with a lower threshold to ignition of a population burst compared to an un-confined culture. This occurred despite the scaling up of the strength of inhibitory synapses. In cultures where density of neurons (as opposed to size of the network, as in our data) was varied, scaling of excitatory and inhibitory synapses also occurred, with excitatory/inhibitory balance preserved for all densities^30^.

**Figure 1.**
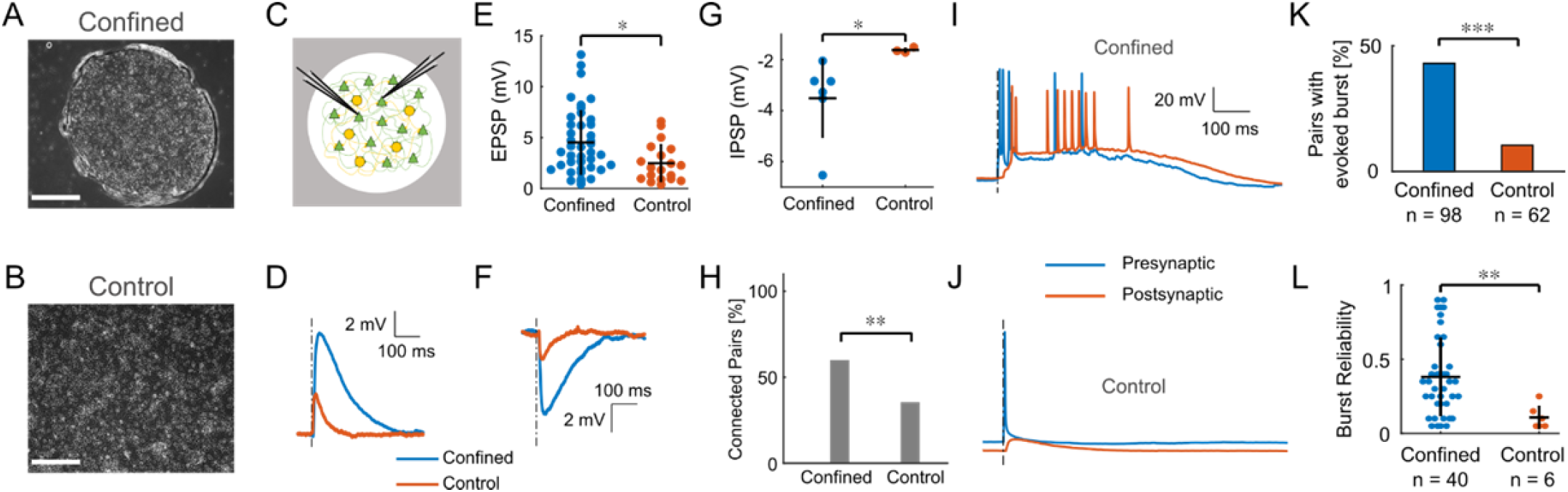
Scaling of excitatory and inhibitory synapses in confined and control networks of cortical neurons. (**A**) Phase contrast images of confined culture and (**B**) control culture. Confinement was provided by a well cut in a polydimethylsiloxane film, which was then attached to the culture dish. Neurons were then cultured inside the well. Scale bars represent 300 µm. (**C**) Dual whole cell recording experiment to record EPSPs and IPSPs. (**D**) Representative EPSPs recorded from neuron pairs in control and confined cultures. (**E**) Peak EPSP amplitudes. (**F**) Representative IPSPs. (**G**) Peak IPSP amplitudes. \**p* = 0.01 and \**p* = 0.02, Wilcoxon rank-sum test (E and G, respectively). (**H**) The percentage of recorded pairs with a functional monosynaptic connection (** p = 0.01, Fisher’s exact test). (**I**) Representative example of evoked population burst in confined culture (burst is evoked in both recorded neurons and in the rest of the network) by stimulating a single neuron. (**J**) Evoked EPSP, but no burst, in neuron pair in a control culture. (**K**) Percentage of neuron pairs in which at least one population burst could be evoked. *** p < 0.001, χ^2^ test. (**L**) Burst reliability in pairs with at least one burst (in response to 10 stimuli). ** p < 0.01, Wilcoxon rank-sum test.

### Hyperexcitability in rate-based computational model

To understand heightened excitability of confined networks, we carried out computer simulations of ring-shaped networks of rate-based neurons. We built networks containing different numbers of neurons, but with the same number of pre-synaptic excitatory and inhibitory connections per neuron (Fig. 2(A)). This resulted in the scaling of synaptic weights with network size – smaller networks had higher average synaptic weights than larger networks (Fig. 2(B)), and higher probability of a synapse existing between a pair of neurons (Fig. 2(C)) (*n* = 3 networks of each size). These results were in line with the findings from dissociated cortical cultures. We then stimulated different numbers of excitatory neurons and allowed network activity to stabilize. We found that average firing rate either went back to zero, or reached maximum rate. We termed the latter ‘population ignition’, as all neurons in the network were firing at maximum firing rate (Fig. 2 (D)). The rate-based model does not include mechanisms for termination of ‘population ignition’, such as afterhyperpolarization or short-term synaptic depression, so the ‘population ignition’ was permanent once it occurred. We did not include these mechanisms since the focus of this study is on seizure initiation. We then determined the minimum number of neurons that needed to be stimulated to achieve population ignition in networks of different size (Fig. 2(E), *n* = 7 networks per size). The relationship between the number of neurons required for ignition and network size was linear, with Pearson *r*^2^ = 0.9985. We examined network dynamics by plotting evolution of excitatory and inhibitory firing rates (trajectories) for different numbers of stimulated neurons. We found that in cases where population ignition was not achieved, inhibitory firing rate increased more rapidly than the excitatory firing rate (Fig. 2(F)). However, stimulation of a larger number of excitatory neurons launched the network on a different trajectory, with excitatory firing rate increasing much more rapidly due to the higher total excitatory synaptic input vs. total inhibitory input to neurons. We found that phase portraits of the 100/10 and 1000/10 networks were similar (Fig. 2 – figure supplement 1(A)), signifying that firing trajectories were determined by the average firing rates. This helps to explain the linear relationship between network size and the minimum number of neurons required for population ignition, as the trajectory depends on initial average excitatory firing rate.

**Figure 2.**
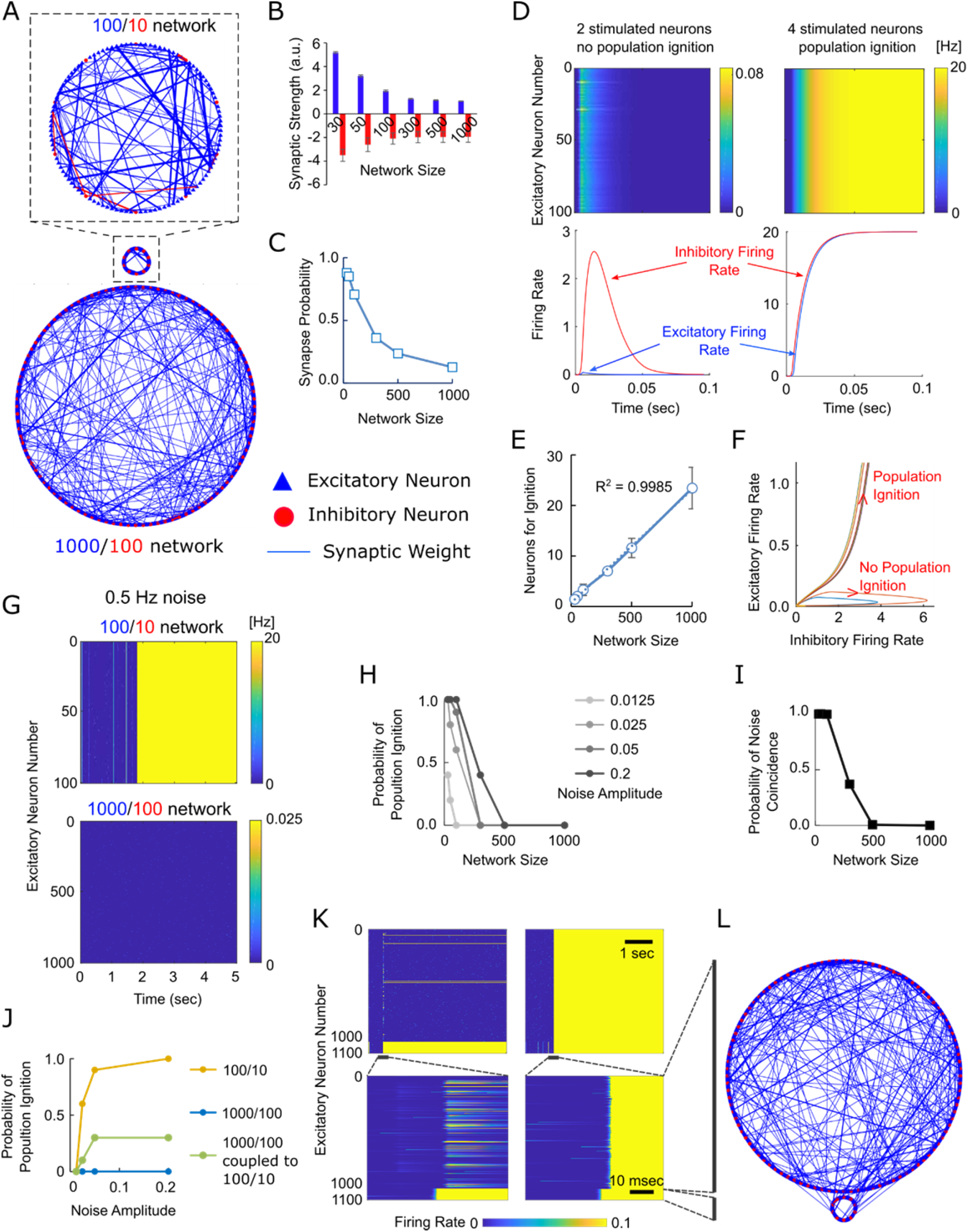
Rate-based model. (**A**) Examples of ring-shaped networks of different size (radius is proportional to the number of neurons in the network). Network with 100 excitatory and 10 inhibitory neurons is 100/10 network; similarly, network with 1000 excitatory and 100 inhibitory neurons is a 1000/100 network. One out of 50 pre-synaptic connections is shown for every 2nd neuron for visibility. (**B**) Scaling of synaptic strength (weight) with increasing network size (specified as the number of excitatory neurons). Excitatory synapses are shown in blue, inhibitory synapses in red. Data are shown as average +/-standard deviation. (**C**) Decrease of probability of an excitatory synapse existing between any two neurons versus network size. (**D**) Representative results of stimulation of 2 or 4 excitatory neurons in a 100/10 network. Top row: raster plots of activity of all excitatory neurons during 0.1 second simulations. Bottom row: plots of average excitatory and inhibitory firing rates versus time, corresponding to raster plots in the top row. (**E**) Minimum number of excitatory neurons that need to be stimulated to achieve population ignition is shown relative to network size. Data points are presented as average +/-standard deviation. (**F**) Representative system trajectories for a 100/10 network where 1 to 10 neurons were stimulated (different color lines correspond to different number of stimulated neurons). (**G**) Representative raster plots for 100/10 and 1000/100 networks where each neuron was stimulated with 0.5 Hz Poisson noise events during a 5 second long simulation run. (**H**) Probability of population ignition in networks of different size when stimulated with noise events of different amplitude. (**I**) Probability of a minimum number of noise events required to ignite the population (I) occurring within a τ_m_ window during a 5 second long simulation, for networks of different size. (**J**) Probability of a population ignition in 100/10, 1000/100, and 1000/100 sub-network coupled to 100/100 sub-network. (**K**) Representative raster plots of a failed ignition (left), and a successful ignition (right) of the larger network (excitatory neurons 1 – 1000) by the smaller network (excitatory neurons 1000 – 1100). Bottom row shows time detail of plots in the top row. Maximum firing rate in these plots is 0.1 to highlight noise events and ignition. (**L**) Representative coupled network of 1000/100 and 100/10 neurons.

We then examined the behavior of networks of different size when stimulated with noise modeled as a Poisson process with events occurring at an average frequency of 0.5 Hz (Fig. 2 (G)). We found that the probability of the population ignition for smaller networks was higher than for large networks, with 1000/100 network having zero population ignition events (Fig. 2(H), *n* = 10 simulations, each network size). These results could be explained by the probability of noise coincidence within a relevant integration window, such as membrane time constant τ_m_. Probability of *k* occurrences in a unit time for a Poisson process is given by:

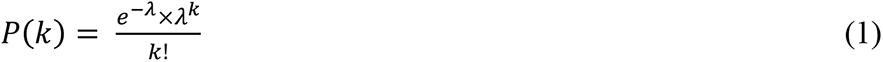

where *λ* is the average occurrence rate per unit time.

In our model,

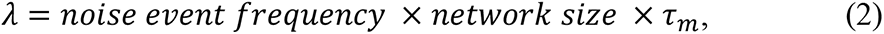

with noise event frequency = 0.5 Hz and τ_m_ = 10 msec. Probability distribution P(*k*) for different values of network size is plotted in Fig. 2 – figure supplement 1(B). We then evaluated the probability that minimum number of neurons required for population ignition will be stimulated by noise event coincidence in 5 seconds / τ_m_ attempts, and plotted the results for networks of different size in Fig. 2 (I). Close correspondence of Fig. 2 (H) and Fig. 2(I) suggests that higher probability of reaching ignition threshold with noise in smaller networks results in higher probability of population ignition, compared to larger networks.

We then examined coupled networks, where 100/10 and 1000/100 networks were connected by replacing some intra-network synapses with inter-network synapses (Fig. 2(L)). We hypothesized that the 100/10 network could act as an ‘igniter’ for the 1000/100 network, and increase the chance of population ignition by noise. We found that this indeed was the case (Fig. 2(J), n = 10 simulations per network size per noise amplitude, with 0.5 Hz Poisson ‘noise’ events occurring in each excitatory neuron). In the case of a coupled network, population was considered to be ignited only if all neurons in 100/10 and 1000/100 portions of the network fired at maximum rate. Examples of a failed ignition (population ignition in the 100/10, but not in 1000/100 sub-network) and a successful ignition (population ignition in both portions of the network) are shown in Fig. 2(K). Raster plots at expanded time scale show that 100/10 sub-network (excitatory neurons 1000-1100) ignited first, followed by ignition in the 1000/100 sub network (excitatory neurons 0 – 1000). Fig. 2 – figure supplement 1(C) shows population ignition with full firing rate scale of 0 to 20 Hz.

### Hyperexcitability due axonal sprouting in spiking computational model

We then investigated whether axon sprouting after injury can result in a locally hyperconnected region that acts as an ‘igniter’ for the larger network. We placed 2000 spiking neurons equidistantly on the surface of a sphere. In this geometry, all neurons have the same number of neighbors. Similar to the rate-based model, 10% of neurons were inhibitory^31^. Excitatory neurons have a total pre-synaptic weight of 300, while inhibitory neurons have a total pre-synaptic weight of 140^27, 32, 33^, with minimum weight of individual synapse equal to 1. For simulation of network activity, synaptic conductance was calculated by multiplying synaptic weight by **g*_syn_*(see Methods). Neurons make their connections stochastically with probability weighted by distance and number of existing connections. Multiple connections between same neurons incremented the synaptic weight by 1. After the intact sphere model matured (neurons reached their connectivity set-point), a localized lesion was modeled by removing a portion of the sphere containing 5%, 25%, 50%, 75%, or 95% of total neurons. After removing neurons, their connections and connections passing through the removed portion of the sphere, different rewiring algorithms were used. Neurons with reduced connectivity compensated by making new connections, modeling structural homeostatic drive. Three different sigmoid distributions of connection probability vs distance were used to model recovery: i) short critical distance, ii) intermediate critical distance, iii) critical distance same as during development (long) (Fig. 3 (A, B)). Re-wiring was terminated after total pre-synaptic weight reached pre-lesion set point. Re-wiring resulted in increased average pre-synaptic weights in the perilesional region (Fig. 3(B)). Inhibitory/excitatory synaptic ratio was not affected by homeostatically driven re-wiring (Fig. 3 (C)). Eight neurons were selected at the north pole of an intact sphere or the edge of a lesioned sphere and the south pole. Tight network clustering^34^ in the microregion at the lesioned sphere edge was observed, contrasting with the loose network clustering in equivalent microregion in the intact sphere, or at the south pole (Fig. 3(D)).

**Figure 3.**
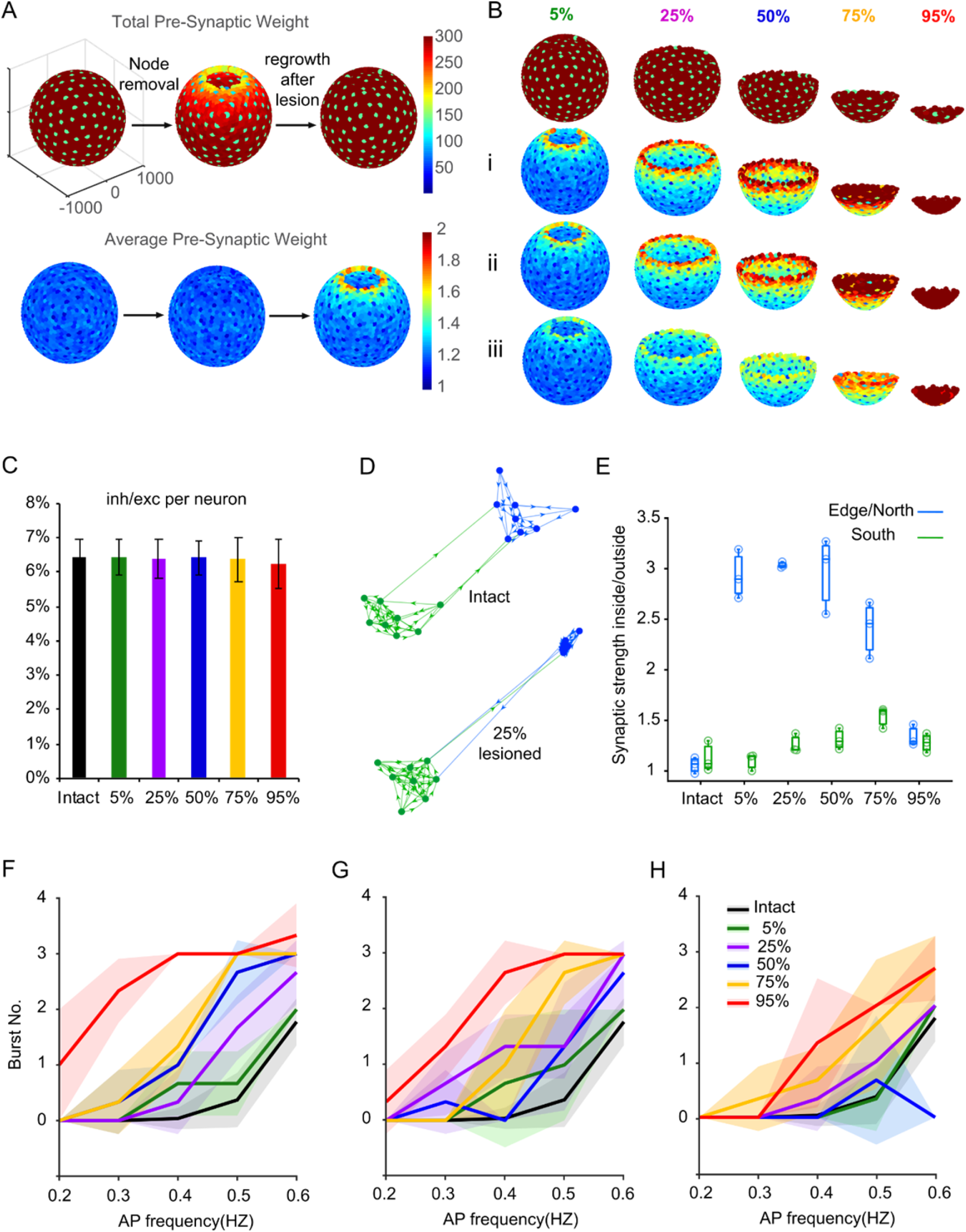
Effect of network size and homeostatically-driven rewiring on synaptic strength and hyperexcitability. (**A**) Fully developed spherical network (left). A portion of the sphere containing 100 neurons (5% of total number of neurons) was removed (middle) to model an injury, and the network was allowed to recover by re-wiring (right). (**B**) Top row shows the total pre-synaptic weight per neuron after re-wiring post injury of different size. Average synaptic weights after re-wiring under different sprouting constraint: (i) Short, (ii) Intermediate, and (iii) Long (developmental) critical sprouting distance. (**C**) Ratio of total inhibitory and excitatory synaptic weight per neuron after injury. No significant difference between groups. (**D**) Representative force-directed visualization of synaptic strengths between neurons at the south pole of the sphere (green), north of the intact sphere (blue, top) or in a micro-region proximal to the cut edge of the 25% injured (lesioned) network (blue, bottom). Stronger synapses are represented as shorter distances between neurons (nodes). (**E**) The ratio of the weight of connections within the blue area in (D) vs. connections from blue to green area in (D), *n* = 3 networks per condition. (**F – H**). Number of population bursts in networks stimulated by Poisson action potentials (APs) at indicated frequency. (F) short, (G) intermediate, and (H) long (developmental) sprouting constraint. *N* = 3 networks per condition, average +/-standard deviation. *P* values obtained with two-sided t-test are provided in Supplementary Tables 1-3.

Connectivity within the microregion near the lesion was significantly higher than connectivity between the microregion and the rest of the network, for all lesion sizes except 95% (Fig. 3(E), *n* = 3 networks generated with different random seeds). No difference of in/out connectivity was observed for equivalent microregion in the intact network (*n* = 3). Lack of difference in connectivity after 95% lesion is due to the small size of the remaining network, with largest distance between any two neurons comparable to critical re-wiring distance.

We then examined the functional outcome of different lesion sizes and re-wiring constraints. Networks were stimulated by Poisson trains of action potentials in individual neurons. Increase in lesion size (and thus decrease in the size of surviving network) resulted in increase in the number of network-wide bursts, indicating increase in hyperexcitability (Fig. 3 (F-H)). Re-wiring with the shortest critical distance resulted in more bursts (Fig. 3(F)) than re-wiring with relaxed critical distance (Fig. 3(G, H)). *P*-values are reported in Fig. 3 – figure supplements 1-3.

### Population bursts initiate in the perilesional region after re-wiring

We explored the effects of rewiring in the perilesional portion of the network on the initiation of population bursts. Out of 14 population events (bursts) that spontaneously occurred in a network with 25% loss, 13 events initiated in the perilesional region. In contrast, event initiation was evenly distributed in the intact network (11 events) (Figure 4 (A-D)). We have also observed focused bursting in the perilesional region (Fig. 4 (E, F)). These sub-population events may represent failed ignitions, similar to those observed in the coupled rate-based network (Fig. 2(K)).

**Figure 4.**
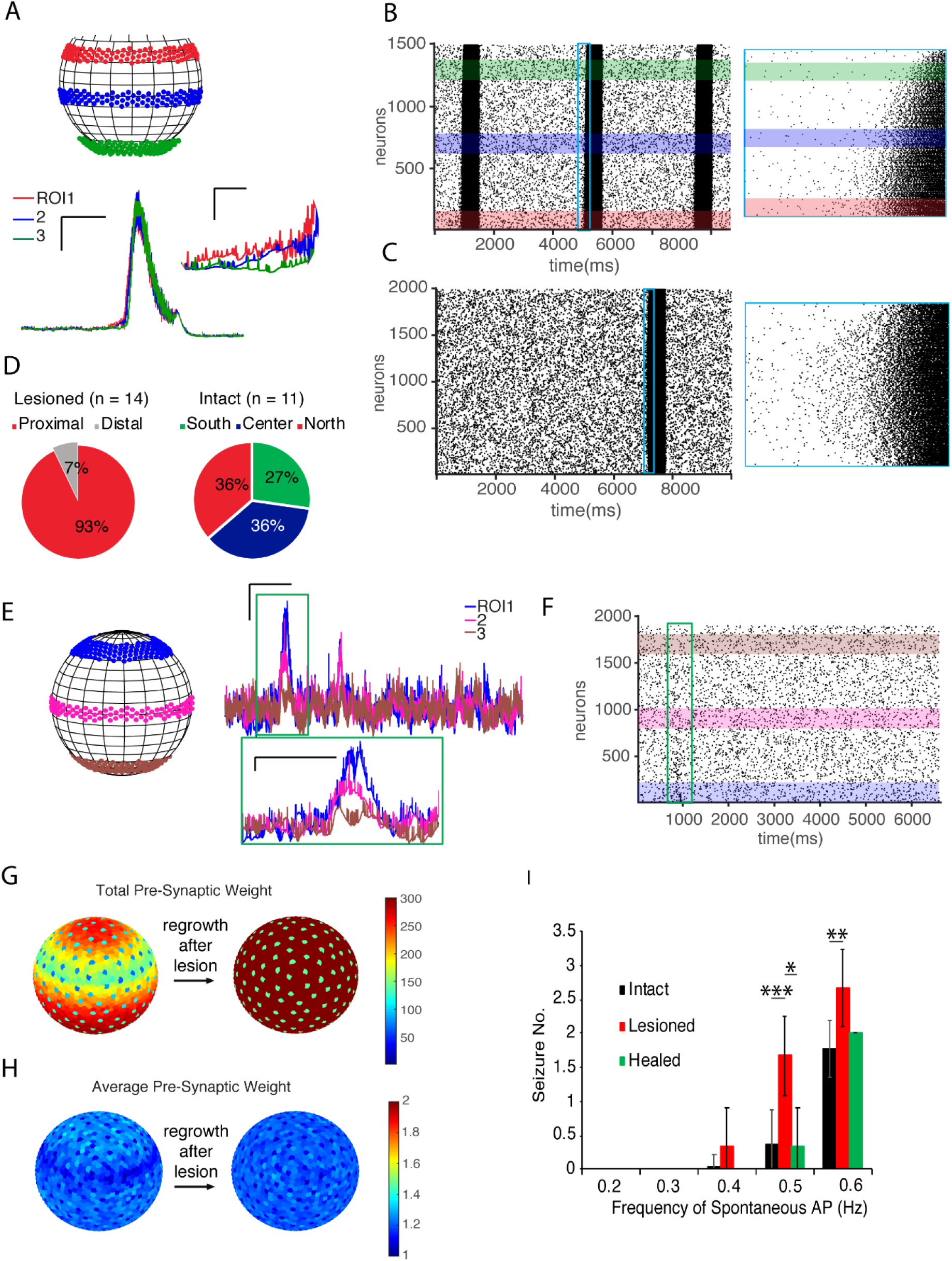
Population burst initiation and network ‘healing’. (**A**) Summed membrane voltages of neurons in regions of interest (ROIs). Scale bars (horizontal, vertical) : 500 ms, 1000 au., inset : 50 ms, 500 au. (**B**) Representative raster plot of spontaneous activity in a lesioned network. Neurons are numbered in order of their distance from the north pole. Right: inset shows initiation of a population burst at expanded time scale (420 ms). (**C**) Representative raster plot of spontaneous activity in intact network. (**D**) Population burst initiation. (**E**) Sum of membrane voltages of neurons in indicated ROIs.for a sub-population event. Scale bars: horizontal, 200 ms, vertical, 500 au. (**F**) Raster plot of activity in (E). (**G**) Total, and (**H**) Average pre-synaptic weight in ‘healed’ networks. (**I**) The number of synchronized bursts (seizures) in intact, lesioned, and healed networks. * p < 0.05, ** p< 0.01, and *** p < 0.001 (t-test).

### Healing the hyperexcitability at the lesion edge by replacing lesioned neurons

We then investigated whether it was possible to rescue (heal) the effect of re-wiring on the network in the perilesional region. We removed all the connections to the lesioned area, but left the neurons in place (Fig. 4 (G, H)). This provided neurons in the perilesional region with additional targets for re-wiring (under short sprouting constraint). The average synaptic weight post sprouting remained comparable to the rest of the network and no strong edge effect was developed (Fig. 4 (G, H)). Excitability of the ‘healed’ network after re-wiring was significantly lower than lesioned network and was not significantly different from the intact network (Fig. 4 (I)), (*n* = 3 networks, each condition).

### Smaller network size correlates with hyperexcitability of organotypic cultures

We created organotypic cultures of different size to experimentally validate findings in computational models (Fig. 5 (A-D)). The number of neurons in these cultures spanned two orders of magnitude, from ∼1000 neurons in the smallest to ∼130,000 neurons in largest cultures (Fig. 5 (E, F)). The number of ictal-like events was inversely related to culture size, with smallest cultures having significantly more ictal-like events than the largest cultures (Fig. 5(G)).

**Figure 5.**
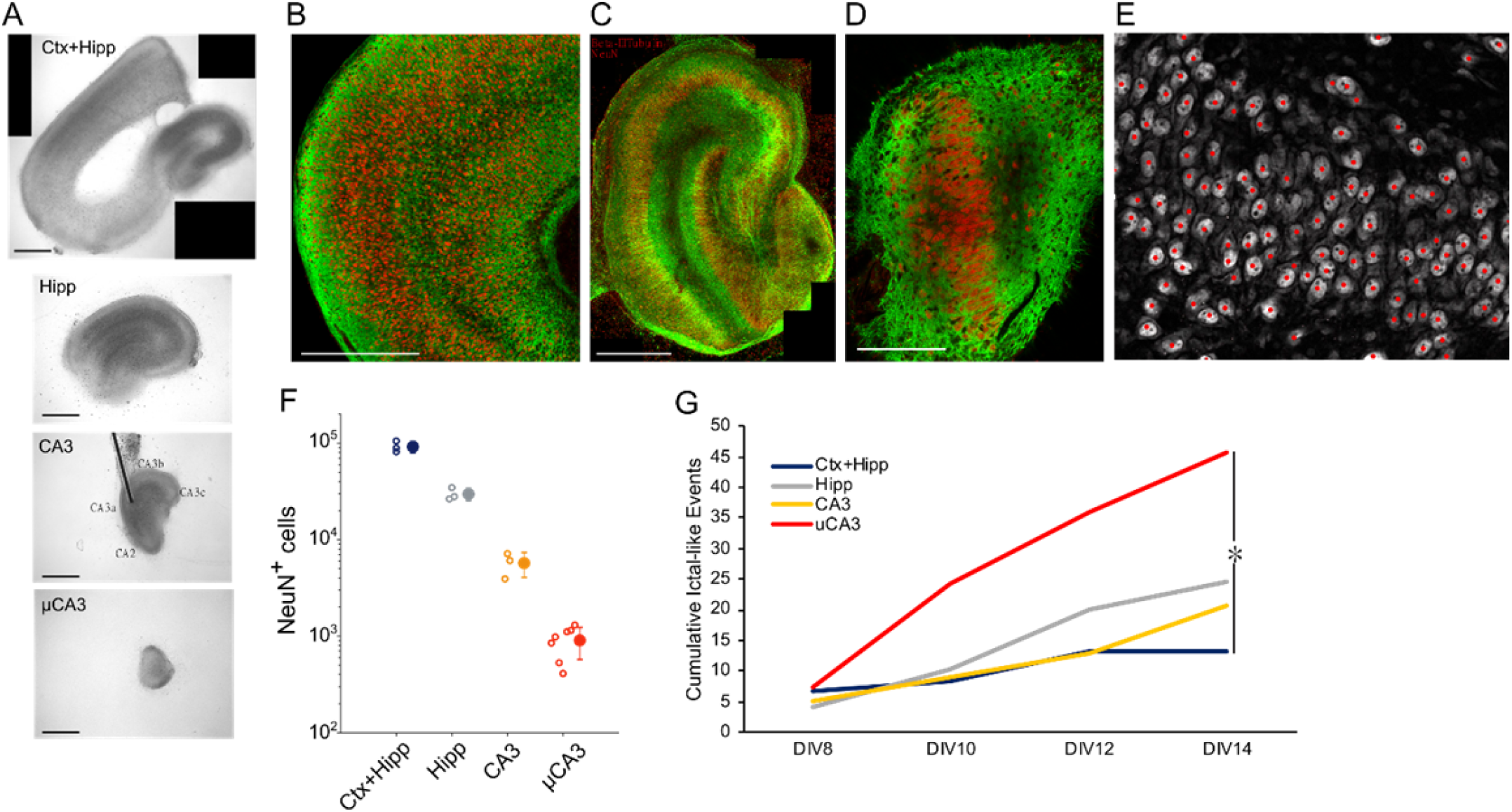
Effect of network size on ictal event occurrence. (**A**) Brightfield images of Ctx+Hipp, hippocampal, CA3 sub-region, and µCA3 (containing a portion of CA3) cultures. (**B-D**) Confocal images from cortical part of a Ctx+hipp culture (B), hippocampal culture (C), and µCA3 (D) culture. Cultures were fixed on 8 DIV and stained for anti-NeuN (red, neuronal nuclei), and beta-III tubulin (green, dendrites and axons). (**E**) Automated neuronal count in 3D optical stack (ImageJ). Counted neuronal nuclei are marked with red dots. (**F**) Number of neurons in the entire culture of each type. *n* (Ctx+Hipp) = 3, *n* (Hipp) = 3, *n* (CA3) = 3, and *n* (µCA3) = 7. (**G**) Cumulative number of ictal-like events in 8 Ctx+Hipp, 25 Hipp, 11 CA3, and 6 µCA3 cultures, KS test *p* < 0.05. Scale bars: (A-C) 500 µm, (D) 100 µm.

### Axon sprouting and functional synaptogenesis in organotypic cultures

We tracked the growth of axons sprouted by neurons in whole hippocampal cultures. We found that axons continued growing rapidly from DIV 2 to DIV 5 (Fig. 5 - figure supplement 1 (A, B)). We did not quantify axon growth at later DIVs, but axons continued growing until at least DIV 14. In order to assess whether sprouted axons can form functional connections, two organotypic slices were cultured together. Slices were placed at distances ranging from 0.5 mm to 1.7 mm from each other. Cultures placed close to each other showed synchronization of ictal-like activity as early as 4 DIV. Cultures at distances of 0.5 – 1 mm showed synchronization by 6 to 14 DIV (Fig. 5 – figure supplement 1 (C, D)). Cultures at distances larger than 1.5 mm failed to synchronize. These findings show extensive axon sprouting by neurons in organotypic cultures that leads to functional re-wiring.

### Clustering of the network in the perilesional region

We next examined whether axon sprouting results in local hyperconnectivity in organotypic hippocampal cultures, as predicted by the computational model (Fig. 3). Injury was experimentally simulated by cutting off different portions of a cultured slice (Fig. 6 (A)), mimicking computational model (Fig. 3 (A, B)). After allowing the slices to re-wire for at least 8 days, we examined the network in the microregion immediately next to the cut (perilesional region) and the larger sub-regional network (Fig. 6 (A)). We found higher connectivity in the microregion near the cut compared to an equivalent microregion in a control culture (Fig. 6 (B)), for cuts delivered to CA3c, CA1, Subiculum (Sub), or cortex (Ctx). Clustering of the network^34^ in the microregion significantly increased in cut versus control cultures (Fig. 6(C), *n* = 8 including 2 technical replicates for 4 cultures, each group). Two-sided Student’s t-test *p* values were *p* = 0.0201 for CA3c-DG, *p* = 0.0024 for CA3c-CA3b, *p* = 0.0042 for CA1, *p* = 0.0016 for Sub, and *p* = 0.0121 for Ctx. Network clustering indicates strengthening of connectivity within the perilesional region, confirming computational results (Fig. 3 (D, E)).

**Figure 6.**
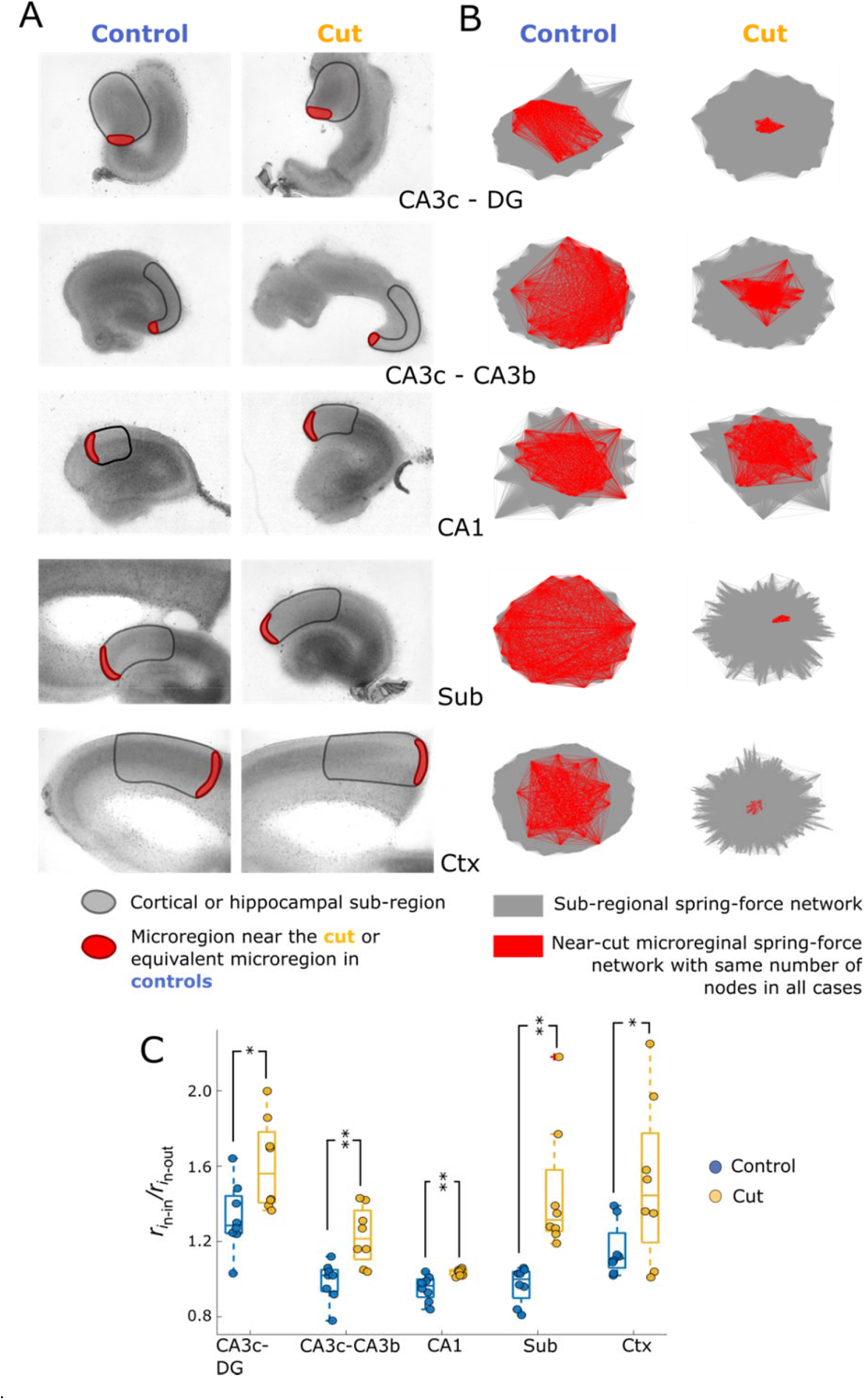
Network hyperconnectivity near the lesion (cut). (**A**) Phase micrographs of control and cut cultures. A sub-region with black outline was selected for correlation-based network analysis. A microregion (red) immediately next to the cut, or an equivalent microregion in control cultures, was selected to determine the level of cut-induced hyperconnectivity. (**B**) Network graphs produced by correlation-based network analysis. Force-directed visualization was used to position nodes so that nodes connected by an edge with large weight experienced a correspondingly strong attraction. Nodes in the microregion and edges between them were highlighted in red. All microregions contained 45 nodes. Tight node clustering indicates strong connectivity between the nodes. (**C**) Ratio of the correlation of nodes in the microregion to each other (*r*_in-in_) and to nodes in the larger network (outlined in black in (A), *r*_in-out_). * *p* < 0.05, ** *p* < 0.01

### ‘Healing’ reversed perilesional clustering

After delivery of a cut to CA3c, the cut edges of the slice were placed into close proximity with each other. Activity in CA3 and DG in the ‘healed’ slice was significantly more correlated than in the cut slice (Fig. 7 - figure supplement 1), *p* = 0.002 (*n* = 7 cut cultures and *n* = 6 healed cultures). This suggested that ‘healed’ cultures re-established synaptic connectivity that was severed by the cut. Network clustering in the ‘healed’ cultures was significantly lower than clustering in ‘cut’ cultures (*p* = 0.0046 for CA3c-DG, and *p* = 0.0049 for CA3c-CA3b), but not significantly different than clustering in controls (*p* = 0.7711 for CA3c-DG, and *p* = 0.1457 for CA3c-CA3b) (Fig. 7). *N* = 8, including 2 technical replicates for 4 cultures were used for each group. Relaxation of clustering suggests that ‘healing’ reduced hyperconnectivity in the perilesional region, similar to computational results in Fig. 4(H).

**Figure 7.**
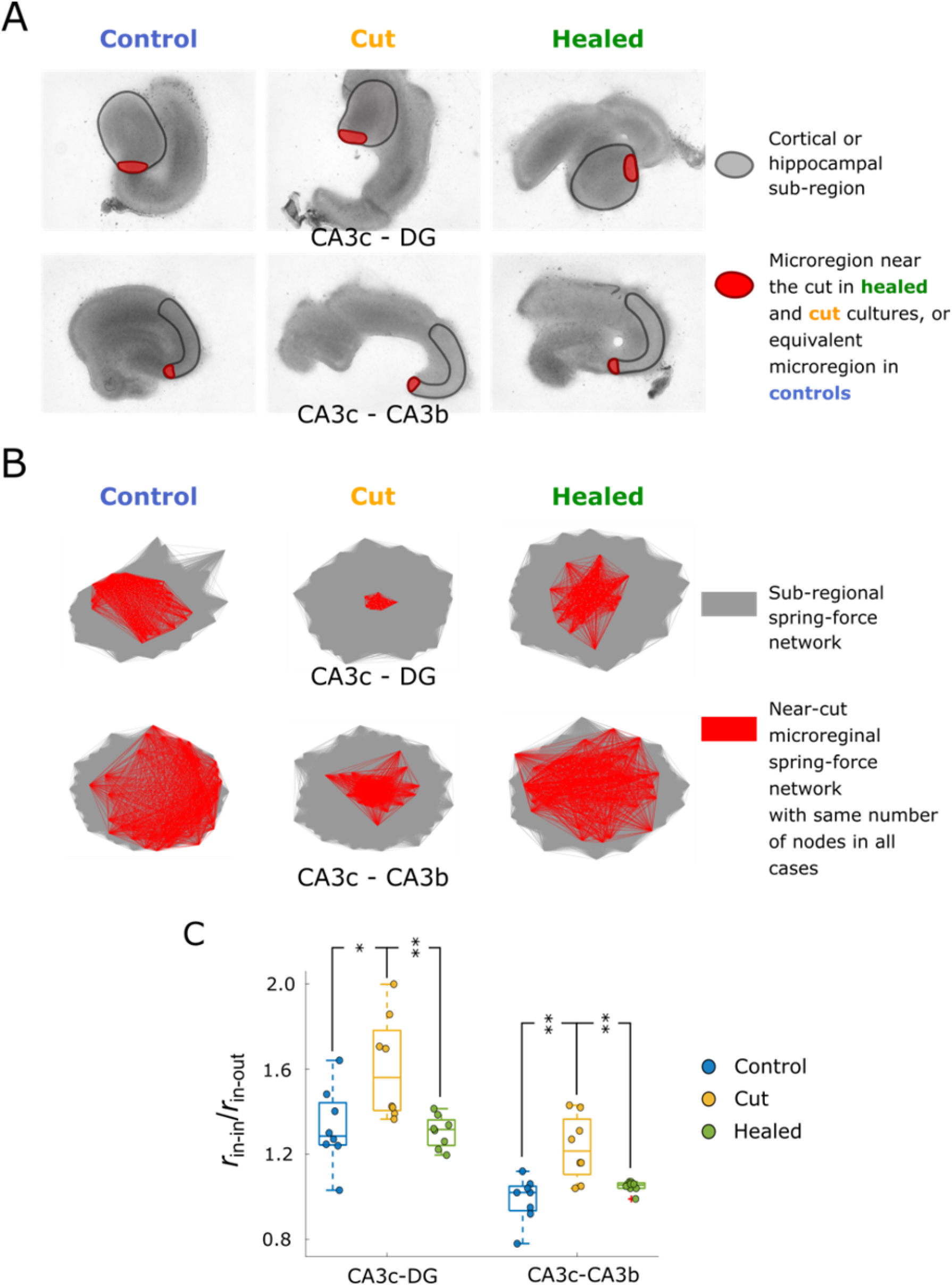
‘Healing’ reverses network clustering. (**A**) Cut was delivered to CA3c. Sub-regions indicated by black outline, and a microregion (red) immediately next to the cut, or an equivalent microregion in control cultures, were selected for network clustering analysis. Control and cut cultures are same as used in Fig. 5. (**B**) Representative force-directed visualizations of network graphs. Nodes in the microregion and edges between them were highlighted in red. All microregions contained 45 nodes. (**C**) Clustering in the microregions in control, cut, and healed cultures. * p < 0.05, ** p < 0.01

### Epileptiform activity initiates in perilesional region

We then examined the effect of cut-induced network reorganization on excitability. We found that perilesional region (near cut delivered to Ctx or subiculum) was predominantly the initiation site for slice-wide ictal-like events (Fig. 8(A-C)). In cultures with cuts at both subiculum and CA3, ictal-like activity initiated in both perilesional regions. ‘Healing’ the CA3 cut abolished ictal initiation near CA3, with ictal events predominantly initiating in subiculum (Fig. 8 (A-C)). Z-scores and corresponding *p* values and sample numbers are shown in Fig. 8 – figure supplement 1. We also examined population events that did not propagate to the entire culture (interictal-like activity, Fig. 8(D)). Energy of interictal activity was significantly higher in the perilesional regions (Fig. 8 (E-H), *n*(ctx+hipp) = 28, *n*(hipp) = 21, *n*(lesioned) = 16, *n*(healed) = 4), suggesting that perilesional region may act as an ‘igniter’ for the rest of the network (Fig. 2 (K, L)).

**Figure 8.**
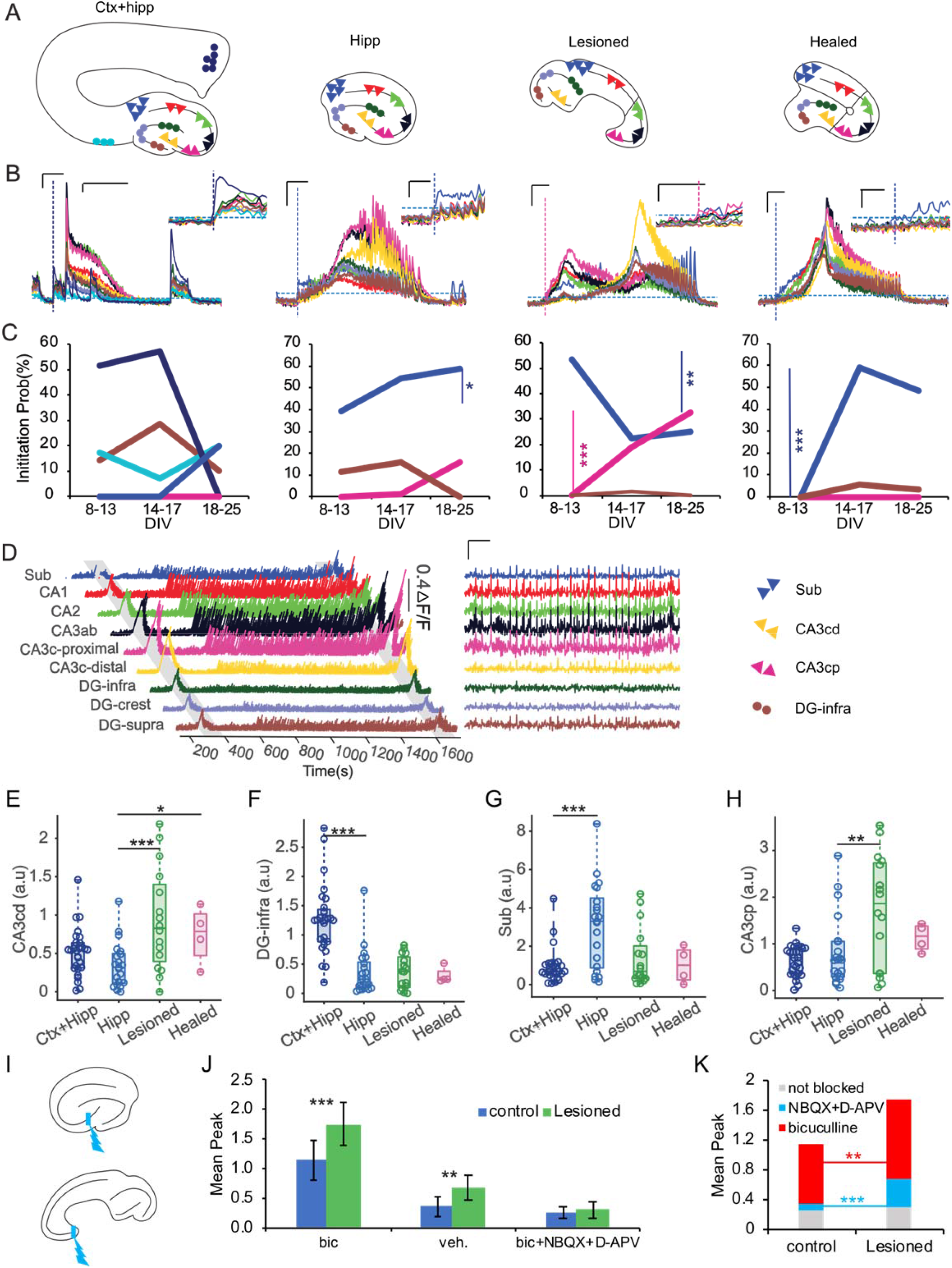
Ictal-like event onset zone in organotypic cultures. (**A**) Ctx+hipp, hippocampal, lesioned and healed cultures. In each culture the ROIs are represented as colored clusters of neurons. (**B**) Representative optical traces of an ictal event for corresponding cultures in (A). Inset shows event initiation. Scale bars: 20 sec, 0.2 ΔF/F, and in the insets: 1 sec, 0.1 ΔF/F. (**C**) Initiation probability in subiculum (dark blue), CA3c-proximal (pink), and DG-supra (brown) for all cultures are compared at different days in-vitro, and cortex-proximal and distal (in light blue and navy blue) are also compared within Ctx+hipp cultures. The probability of microregion closest to the cut initiating seizures increases with time in vitro, probably reflecting increased connectivity. In healed cultures, no participation of CA3c-proximal in seizure generation was observed. Significance of population proportions was assessed through z-scores. (**D**) Left: example of population activity in a culture with lesion in CA3. Two ictal events are shaded. Right: zoomed-in view of inter-ictal period (scale bars : 10 sec, 0.1 ΔF/F). Energy of inter-ictal activity in CA3ab and CA3c-proximal subregions are significantly stronger than in other subregions. (**E-H**) Energy of inter-ictal activity. (**I**) Microregion near the lesion and comparable microregion in a control culture were optically stimulated as shown, and evoked response from region indicated with blue line measured. (**J**) Peak evoked responses in vehicle, bicuculline, and bicuculline + D-AP5/NBQX. (**K**) Differences of average peak responses measured in drug and in vehicle were calculated, \**p* < 0.05, \*\**p* < 0.01, and \*\*\**p* < 0.0001 (two-sided t-test).

### Perilesional region has increased excitatory and inhibitory synaptic drive

We delivered optical stimulation to the perilesional region and an equivalent region in control cultures (Fig. 8(I)). Evoked activity in the perilesional region was significantly higher compared to control. Significant difference persisted after application of bicuculline, but was abolished after application of bicuculline together with NBQX and D-APV (Fig. 8(J)). Biological replicates were as follows: vehicle control *n* = 5(5) and lesioned *n* = 9(5), bicuculline and bicuculline + D-AP5 + NBQX control: *n* = 11 (6) and lesioned *n* = 11(6), where *n* = recordings(cultures). Increase in evoked response due to bicuculline was significantly higher in perilesional region compared to control, while decrease in evoked response due NBQX and D-APV was also significantly higher in perilesional region compared to control (Fig. 8(K), *p* = 5.776*10^-4^, and 0.009, t-test performed on *n* = 50 randomly drawn differences (bic – veh, and veh – (NBQX+D-APV)) from data in Fig. 8(J)). This suggests that excitatory and inhibitory synaptic drive was higher in the perilesional region.

## Discussion

Organotypic hippocampal slice cultures are a model of post-injury epileptogenesis, where injury is delivered by isolation of the hippocampal slice^35^. Surviving neurons in the slice sprout axons and become hyperconnected over time, with robust excitatory and inhibitory synapses^36^. The resulting network is hyperexcitable, and epileptiform activity emerges after a latent period^37^. Slice cultures thus contain a re-wiring network where the availability of synaptic partners is limited by network size. Homeostatic control of sprouting can be expected to result in increased probability of connections and increased synaptic strength between neurons in smaller networks. Our computational modeling showed that networks of different size, but with same excitatory/inhibitory synaptic drive, require the same proportion of the excitatory neurons to fire to ignite a population event. A direct consequence is that in smaller networks, firing of fewer excitatory neurons is required to ignite the rest of the network. Higher statistical likelihood of noise coincidence in a small network in turn leads to a higher probability of population events. This theoretical/computational result is supported by our finding that cultures containing fewer neurons have more ictal-like events.

Axon sprouting and network reorganization in slice cultures are not homogeneous. A hippocampal slice is disconnected from the surrounding tissue at its horizontal top and bottom planes, as well as at the transverse cut through subiculum. Surviving neurons near the cut are the most disconnected. Ability of these neurons to establish long-distance connectivity is limited by absence of normal synaptic partners and developmental axonal guidance cues. Homeostatic connectivity drive and inability to form long-distance connections can be expected to result in a hyperconnected local network in the region near the transverse cut (perilesional region). We observed this experimentally, and found that formation of hyperconnected local network occurred near the cut delivered to subiculum, entorhinal cortex, CA1, or CA3. Hyperconnectivity was abolished by restoring the number of available synaptic partners. This ‘healing’ experiment also demonstrated that hyperexcitability of the local network was not due to non-network effects of the cut such as inflammation or changes in intrinsic neuronal excitability. Increases in excitatory and inhibitory responses to stimulation are consistent with formation of hyperconnected network in the perilesional region. The local networks generated by the cut in entorhinal cortex, subiculum, CA3, and CA1 all exhibited a higher occurrence of local population events. Ictal-like events were frequently, but unreliably initiated by local activity in the hyperconnected microregion. These experimental results are consistent with the computational result where limitation of the distance of axon sprouting after network lesion led to formation of hyperconnected, seizure initiating local network. Hyperexcitability of the local hyperconnected network may be due to a lower threshold for local network ignition, as described above for small isolated networks. Coupling of small, local hyperconnected network, and a larger network, results in the local network functioning as an unreliable igniter of the larger network. The unreliability may be due to variability in the small network ignition trajectory, which in turn gives a chance for the inhibitory neurons in the larger network to activate faster than excitatory neurons and prevent full network ignition (Fig. 2(K)). This is consistent with the concept of “inhibitory surround” controlling seizure propagation in slices^38^ and animals^39^, and supported by recordings from patients^40^. Increases in inhibitory firing prior to seizure initiation^41^ may represent trajectories of failed ignitions (Fig. 2(D)). The excitatory-inhibitory dynamics of our model are thus in-line with clinical and experimental data, and help explain how sprouting leads to hyperexcitability.

Functional and structural homeostatic mechanisms are involved in network response to injury^42^. In dissociated cortical cultures, the strength of newly formed synapses inversely scales with culture density^30^. This is indicative of a homeostatic process that leads to formation of more synapses with the same partner if fewer partners are available (low density culture), or fewer synapses with the same partner if more partners are available (high density culture). We found that changing the size of dissociated culture, and thus the number of available synaptic partners, also leads to the same effect. The ratio of inhibitory to excitatory synapses was preserved in both low and high density cultures^30^, consistent with our finding that both excitatory and inhibitory synapses scale up in confined networks. Hyperexcitability in confined networks is therefore more likely to be a function of altered network dynamics, rather than altered inhibitory/excitatory balance. One of the mechanisms of homeostasis is preservation of activity level in neurons. Pharmacological suppression of activity leads to an increase in excitatory synaptic conductivity. This functional homeostasis is paralleled by an increase in number of synaptic contacts – structural homeostasis^42^. Injury that results in disruption of neural connectivity may lower neural activity, and thus provide homeostatic impetus for synaptogenesis and strengthening of existing synapses. However, activity can also result in synaptic pruning, and the role of activity in rewiring of inhibitory connections is less clear. We have therefore elected to use excitatory and inhibitory synaptic drive as a homeostatic set point in our computational models. This set point may be the result of neuronal activity levels, or a consequence of homeostasis of neuronal metabolism, synaptic protein synthesis and degradation. Cell signaling kinases that play a role in regulation of metabolism and protein synthesis, such as receptor tyrosine kinases, PI3K, and mTOR, have been considered as antiepileptic targets^43, 44^. Pharmacological inhibition of mTOR resulted in reduction of axon sprouting, suggesting that this kinase may be involved in maintenance of structural homeostasis, potentially through its role in regulating protein translation. However, reduction of axon sprouting did not always prevent epileptogenesis^45^. This may be due to differential role of mTOR inhibition on inhibitory and excitatory sprouting^46^. Our results suggest an alternative interpretation: reduction in the length of sprouted axons may lead to increased local synaptogenesis, which leads to formation of local, hyperconnected network with a lower ignition threshold. This may reduce anti-epileptogenic effects of mTOR inhibition. Interventions that reduce density of local sprouted axonal collaterals may be more beneficial from this point of view.

The goal of epilepsy resection surgery is the removal of a portion of the brain that is seizure-genic. Resection surgery is highly effective, with majority of patients seizure-free one year after the surgery^47^. However, surgery recurrence rate increases with time after surgery. Network reorganization, including strengthening of connections, occurs in patients in the months or years after resection, and can be associated with seizure recurrence^48^. Removal of brain tissue via resection necessarily results in loss of connections by the surviving neurons. Lost connections may include those with synaptic partners in the removed tissue, and those where the connecting axons passing through the removed tissue were severed. This situation is similar to our computational and in vitro models where removal of a portion of a network resulted in loss of connections by surviving neurons. Our results suggest that this resulted in formation of hyperconnected, hyperexcitable local networks in very close proximity to the removed portion of the network. A similar process may occur in patients after surgery, and may be one of the causes of seizure recurrence. The small spatial extent of the hyperconnected networks suggests that development of local treatments may be a promising direction in preventing seizure recurrence after epilepsy surgery.

## Conclusion

Our computational and experimental findings help to explain how post-injury axon sprouting and balanced excitatory/inhibitory synaptogenesis lead to hyperexcitability, and may elucidate one of the mechanisms of the development of epilepsy after injury. This mechanism may also be present in other types of maladaptive neuronal responses to injury.

## Materials and Methods

### Synaptic Scaling in Dissociated Cortical Cultures

Dissociated cortical neurons from post-natal day 0–1 Sprague-Dawley rat pups (Charles River Laboratories) were prepared as described in ^49^. We plated approximately 1 μl of dissociated cell solution (10,000 cells/µl) in well punched in a PDMS film of 100 µm thickness on a poly-D-lysine coated dish. For the first one hour after plating cells, cultures were incubated at 37⁰C and 5% CO_2_ in a medium containing 10% Fetal Bovine Serum (FBS) in Neurobasal-A supplemented with GlutaMAX and gentamicin (Fisher Scientific). After 1 hour, the medium was replaced with fresh culture medium (97.45% Neurobasal-A, 2% B27, 0.25% GlutaMAX and 0.3% gentamicin, all from Fisher Scientific) and the cultures were placed again inside the incubator. Half of the medium was replaced with fresh medium twice a week. All animal use protocols were approved by the Institution Animal Care and Use Committee (IACUC) at Lehigh University and were conducted in accordance with the United States Public Health Service Policy on Humane Care and Use of Laboratory Animals.

For paired whole cell recordings, we replaced the culture medium with an ACSF solution. The ACSF solution contained (in mM): 140 NaCl, 2.4 KCl, 10 HEPES, 10 glucose, 2 CaCl_2_, 1 MgCl_2_, 1 Na_2_HPO_4_ (p H 7.4). Recordings were performed with Patch Pro 1000 (Scientifica) at 37⁰. Electrodes had 5-10 MΩ resistance when filled with internal solution containing (in mM): 130 k-gluconate, 10 HEPES, 10 phosphocreatine, 5 KCl, 1 MgCl_2_, 4 ATP-Mg and 0.3 mM GTP [3]. Cells were visualized through water immersion 40X objective mounted on Axio Examiner microscope (Zeiss) equipped with differential interference contrast (DIC) optics. Multiclamp 700B (Molecular devices) was used for whole cell recordings. Data were acquired at 10 KHz using Digidata 1550B (Molecular Devices). Analysis of recordings was performed in Matlab 2020b (MathWorks, Natick, MA, USA). The presynaptic neuron of each pair was injected with 10 current pulses at 0.1 Hz. Post synaptic potentials were measured as a median of 10 responses.

### Firing Rate – based Computational Model (code available online^50^)

This model is based on firing rate-based (non-spiking) neuronal units with piece-wise linear threshold adopted from^51^. Output of the units is time dependent firing rate *r*:

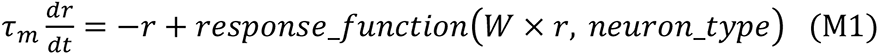

where τ_m_ is membrane time constant, *W* is the matrix of synaptic weights, where weight *W*_ij_ is a positive (excitatory synapse) or negative (inhibitory synapse) integer, and *response_function* for neuron *j* is 0 if input to neuron *j* is < *threshold*, or equal to slope * (input – *threshold*) or maximum firing rate *max_rate*. Slope for *neuron_type* = inhibitory neurons (*inh_slope*) is higher than the slope of the *neuron_type* = excitatory neurons (*exc_slope*) to reflect their higher firing rate and lower spike frequency adaptation.

Neurons are arranged at equal distances along a perimeter of a circle with radius that scales with the number of neurons in the network,

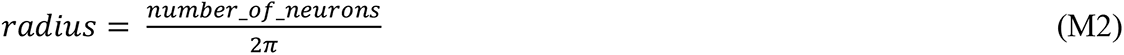

such that distance between two neighboring neurons is the same regardless of network size. Total weight of excitatory synapses (*max_exc_input*) and inhibitory synapses (*max_inh_input*) is the same for all neurons and all networks, regardless of network size. Individual synapses are assigned based on exponentially decaying distance-depended probability (distance is the arc length between any two neurons), with distance constants λ_exc_ and λ_inh_. Synapse assignment is iterated until *max_exc_input* and *max_inh_input* are reached, with individual synaptic weights incremented each time assignment is made to the same synapse.

In the case of the coupled network, two sub-networks are placed at distance equal to half of the radius of the small network, and 1-2% of total excitatory synaptic weight of the smallest sub-network is randomly removed from both sub-networks. Synapses are then replaced with trans-network connections, using the same iterative process as above.

Stimulation was delivered as a single time step increase in rate, while noise was delivered as randomly occurring pulses with width of a single time step, average frequency of 0.5 Hz, and specified amplitude.Simulation parameters are shown in Table 1. The network was simulated using Matlab 2020b (MathWorks, Natick, MA, USA).

**Table 1.** Rate-based model parameters.

| $\tau_m$ (msec) | threshold | exc_slope | inh_slope | max_rate<br>[Hz] | max_exc_input | max_inh_input |
| --- | --- | --- | --- | --- | --- | --- |
| 10 | 0.1 | 0.01 | 0.043 | 20 | 150 | 10 |

#### Pseudocode for single network with stimulated neurons

%Parameter Definition

set inhibitory and excitatory neuron parameters

set network parameters

set stimulation amplitude

set simulation timestep and duration

*%*Create *Network*

Position excitatory and inhibitory neurons around a ring, where radius is proportional to network size

~~~
**While** set excitatory and inhibitory input has not been reached
        **for** each postsynaptic neuron
                Select a set of possible presynaptic neurons
                **for** each selected neuron
                        **if** neuron is excitatory
                                **if** dice roll is < *e*^-distance between neurons / critical excitatory distance^
                                             create connection
                        **else if** neuron is inhibitory
                                **if** dice roll < *e*^-distance between neurons / critical inhibitory distance^
                                             create connection
~~~

set neuron self-connectivity to zero

%*Run Simulation*

~~~
**for** selected number of stimulated neurons
       randomly select set of stimulated neurons
       create stimulation input at a single time step
       **for** each time step
              Neuron input = connectivity matrix * neuronal firing rates
~~~

Determine neuronal firing rate on next time step based on current firing rate, thresholded response to input, and stimulation

#### Pseudocode for coupled network with noise

*%Parameter Definition*

set inhibitory and excitatory neuron parameters

set network parameters

set number of neurons for large network

set number of neurons for small network

set weight of connections between large and small network

set Poisson noise event amplitude and frequency

*%*Create *Large Network*

Same steps as in Single Network Creation

*%*Create *Small Network*

Same steps as in Single Network Creation

*%*Create *Combined, Coupled Network*

~~~
**while** weight of replaced connections < weight of connections between large and small network
        randomly select a neuron and an input synapse
                **if** synapse is excitatory
                        remove the synapse
~~~

randomly select an excitatory neuron in the other network and make a new synapse

increment weight of replaced connections

%*Run Simulation*

create Poisson noise events for each neuron

**for** each time step

~~~
       Neuron input = connectivity matrix * neuronal firing rates
~~~

Determine neuronal firing rate on next time step based on current firing rate, thresholded response to input, and stimulation by Poisson noise

### Spiking computational model of homeostatic sprouting and synaptogenesis after injury (code available online^50^)

2000 nodes are distributed on a sphere with radius of 950 μm, including 1800 excitatory and 200 inhibitory neurons. Therefore 10% of all the neurons are inhibitory^31^. Nodes are positioned equidistantly on the sphere which is described by its cartesian coordinates x, y, and z^52^. The arc distances are calculated by calculating the angle between any two nodes. Each node has a defined number of indegree and outdegree connections as the “set point”. Excitatory neurons receive 300 synapses in total, and inhibitory neurons receives 140 connections. 6-7% of inputs for each neuron are inhibitory^27, 32, 33^. Only fast-spiking inhibitory interneurons making synapses with excitatory cells were considered in this model. Connectivity is developed via multiple iterations. On each iteration, η neurons are randomly selected based on a sigmoidally weighted probability, and η synapses are made on the appropriate postsynaptic neuron. The postsynaptic neuron is also chosen randomly based on a weighted probability. Sigmoid weighting represented distance dependent distribution. It has been reported that bellow 400μm, the probability of connection does not depend on distance^53^; however, the probability decreases exponentially for longer distances ^54, 55^. The distributions are different for excitatory neurons and inhibitory neurons, since the excitatory neurons can extend their axons over much longer distances and inhibitory neurons make more local connections. Moreover, as the neurons get closer to the homeostatic connectivity setpoint, their priority to be picked drops based on a sigmoid distribution. Therefore, the product of the distance dependent distribution and the availability of neuron to receive a new connection determines probability for a neuron to be picked as a postsynaptic partner. After neurons reach the final setpoint, the current state of the network is considered as a fully matured state.

After connectivity was complete, leaky integrate and fire model was used to simulate neurons:

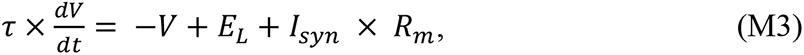

where **τ** is membrane time constant, E_L_ is resting membrane potential, *I*_syn_ is the synaptic current, and R_m_ is the membrane resistance. The following parameters were used:

R_m_(inh) = 400 MΩ, R_m_(exc) = 150 MΩ, E_L_ = −65 mV, **τ*(*in*ℎ)* = 5 ms, **τ*(*exc*) =* 50 ms and dt = 1 ms.

Synaptic current was calculated as follows:

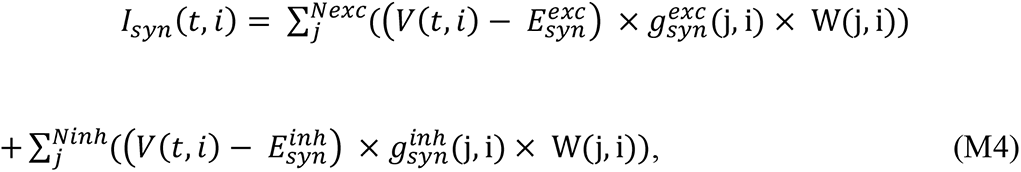

where 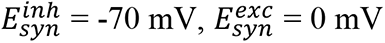, and *N_exc_* and *N_inh_* are the numbers of all the excitatory and inhibitory neurons that have synapses on neuron *i*. Synaptic conductance was implemented by a single exponential function **g*_syn_* = **g*_max_ (*t*) *e*^-t/τsyn^*. Short term depression was implemented by decaying **g*_max_* after every action potential *g*_*max*_(*t* + 1) = *g*_*max*_(*t*) × 0.975, and allowing it to recover when no new action potentials arrived at the presynaptic terminal: If 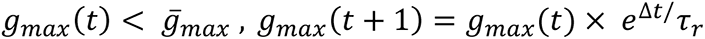, where τ_*r*_ = 1000 msec. Maximum conductances were: 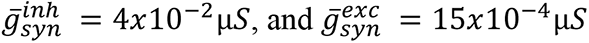.

The time delay from the presynaptic neuron firing an action potential and inducing a postsynaptic potential is calculated based on the arc distance between neuron i and j and is as follows:

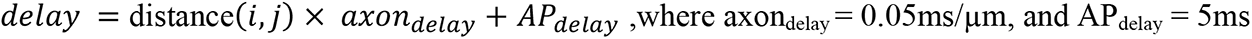

We introduced noise (spontaneous activity) by making each neuron fire action potentials as a Poisson train with specified average frequency. All simulation runs were 10 seconds long.

### Organotypic Cultures

Hippocampi and entorhinal cortices of 7-8 day old Sprague-Dawley rats were dissected and slices with 350 μm thickness were made using McIlwain tissue chopper (Mickle Laboratory Eng. Co., Surrey, United Kingdom). Slices were randomly assigned to experimental groups and cut as described in Results section. Slices were then placed onto poly-D-lysine (PDL, Sigma Aldrich) coated 6-well tissue culture plates or 35 mm tissue culture petri-dishes (Falcon). Slices were maintained in a humidified 37°C incubator with 5% CO2 regulation on a rocking platform. A serum free culture medium consisted of Neurobasal-A/B27, 30 μg/ml gentamicin, and 0.5mM GlutaMAX (Fisher Scientific). Culture media was changed bi-weekly. All animal use protocols were approved by the Institution Animal Care and Use Committee (IACUC) at Lehigh University and were conducted in accordance with the United States Public Health Service Policy on Humane Care and Use of Laboratory Animals.

### Electrophysiological Recordings and Analysis

6-well tissue culture plates were transferred to an interface chamber (Bioscience Tools) connected to a temperature controller maintaining temperature at 37 °C and a blood gas providing 5% CO_2_, 21% Oxygen, balanced Nitrogen (Airgas). Tungsten microwires with 50 μm diameter non-insulated tip were placed under CA3 or CA1 neuronal layers and extracellular field potentials were recorded for 45 minutes via high-impedance multiple-channel pre-amplifier stage (PZ2-64, Tucker Davis Technologies) connected to a RZ2 amplifier (Tucker Davis Technologies). Signals were filtered with a band-pass filter (1 Hz - 3 KHz, gain x1000) and sampled at 6 KHz rate. Recorded signals were analyzed using OpenX software (Tucker Davis Technologies) and MATLAB (MathWorks). The first 15 minutes of each recording were not included in the analysis. A threshold of 65 μV was applied to negate background noise. Seizures (ictal-like events) were identified as paroxysmal events with frequency greater than 2 Hz that lasted for at least 10s.

### Optical Recordings

jRGECO1a was expressed under synapsin promoter in organotypic cultures by applying pAAV.Syn.NES-jRGECO1a.WPRE.SV40 (titer ≥ 1010) to the culture medium on 0 DIV and incubating for 1 hour. Half of the medium was replaced with fresh culture media on 3 DIV and after that whole culture medium was changed bi-weekly. pAAV.Syn.NES-jRGECO1a.WPRE.SV40 was a gift from Douglas Kim & GENIE Project (Addgene plasmid # 100854; http://n2t.net/addgene:100854; RRID:Addgene_100854). Culture dishes were placed into the mini-incubator while keeping the temperature at 37 °C and supplementing with blood gas, on a fluorescent inverted microscope stage (Olympus). Recordings were 15 to 30 minutes long. CCD camera and 4X objective was used to record fluorescent changes at frame rates ranging from 1.59 seconds/frame to 0.16 seconds/frame (∼0.6-6 Hz). Videos were then analyzed using ImageJ and MATLAB.

### Image Processing

Regions of interest were drawn for different cultures and mean gray values were calculated for each frame and recorded in .txt files using ImageJ drawing tools and Plugins. Values were then imported to MATLAB and baseline was calculated for the optical signal using the asymmetric least square smoothing method^56^. Signal to baseline ratio was calculated by the following equation:

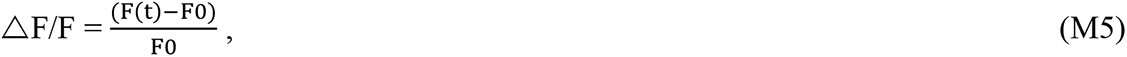

where F0 is the baseline. In order to identify paroxysmal activities, a simple thresholding method was used. The threshold was greater than three times the standard deviation of baseline activity. For calcium intensity analysis, all the data points above the threshold were summed for each region of interest (ROI) and normalized to the activity of the whole culture. For seizure-like event detection we used the same criteria we used for electrical recordings; the optical signal above the threshold must last for at least 10 seconds to be counted as a seizure.

### Axon growth tracking

Slices were cultured in 6-well plates and phase contrast images were taken with a 20X objective from 2 DIV to 5 DIV. Two sets of images were collected on each day 2 to 4 hours apart. Trackable axon terminals were marked by arrows, and the length by which axons grew was measured.

### Correlation-based Network Analysis

Optical recordings (250 msec/frame) of jRGECO1a fluorescence in control, cut, and healed cultures on DIV 8 – 12 were used. Activity was analyzed in a region of interest encompassing hippocampal or cortical sub-region located near the cut, or an equivalent region of interest in control cultures. Region of interest was split into bins of 2×2 or 3×3 pixels, and ΔF/F timeseries of approximately 30 minutes were calculated for each bin. Pearson correlations coefficients *r* were then calculated for each pair of bins. Bins in microregions immediately proximal to the cut, or in equivalent microregions in control cultures, were then selected (45 bins in each microregion in all analyzed cultures). Averages were then calculated for *r* between bins within the microregion (*r*_in-in_), and between bins within microregion and bins outside of microregion(*r*_in-out_). Adjacency matrices for each sub-region were populated with *r* values between appropriate bins. Network based on this adjacency matrix was then plotted using ‘force-directed layout’ option in Matlab graph plot command. Effect of edge weights (*r* coefficients) was selected to be inverse; in other words, largest edge weights produced strongest attractive force and thus closest spacing between nodes (bins). Bins belonging to the microregion, and edges between them, were highlighted in red on the resulting graph plot to show the relationship between sub-regional and microregional networks.

### Detection of ictal initiation zone

Each culture was recorded at least three times between 8 and 25 DIV. Multiple ROIs within each culture type were selected. ROIs considered were: subiculum, CA1, CA2, CA3b, CA3c-proximal, CA3c-distal, DG-infra, DG-crest, DG-supra for hippocampal, lesioned, and healed cultures. Two extra ROIs were considered for Ctx+hipp cultures as well: Ctx-sub and perirhinal-Ctx. We then traced the evolution of calcium activity for each ROI within the culture. A zero-phase low pass filter was applied to the optical signal to remove noise. An event is considered ictal if the optical trace for all the ROIs (generalized events within hippocampus) was above the threshold for more than 10 second without dropping below threshold for more than 0.5 seconds. However, for the Ctx+hipp cultures activity in the perirhinal-Ctx and Ctx+sub ROIs were exempt from this requirement as they are not present in other culture types. In addition, in lesioned cultures ROIs located at CA3c-distal and DG regions were also exempted from the requirement as in some cases they are distally located and act as separate network. Initiation zone probability was calculated as the number of ictal events originating in a given ROI over the total number of ictal events that were evaluated.

### Energy of Interictal Activity

Energy was defined as the area under the curve (a.u.c) of the ΔF/F(t) during interictal period.

### Patterned Optical Stimulation

Cultures were infected with both pAAV.Syn.NES-jRGECO1a.WPRE.SV40 and pAAV-hSyn-hChR2(H134R)-EYFP(AAV9) (titer ≥ 1013) on DIV 0. Channelrhodopsin (ChR2) is a light sensitive ion channel which when expressed localizes into the membrane of the neuron. pAAV-hSyn-hChR2(H134R)-EYFP was a gift from Karl Deisseroth (Addgene plasmid # 26973; http://n2t.net/addgene:26973; RRID:Addgene_26973). The recording media consists of NaCl 134 mM, KCl 2.4 mM, HEPES 10 mM, Glucose 10 mM, CaCl_2_ 2 mM, MgCl_2_ 1 mM, and Na_3_PO_4_ 1.2 mM. Simultaneous optical stimulation of ChR2 and optical recording of jRGECO1a fluorescence were carried out by using a dual-deck inverted fluorescence microscope (Olympus IX73) equipped with X-cite LED illuminator on the lower deck and a digital micromirror device-based patterned illuminator (Mightex Polygon400) on the upper deck. For these experiments, 20X objective was used and light stimuli were delivered via patterned illuminator to a narrow rectangle at the desired location on a cultured slice. Stimulation was delivered as 10 blue light pulses at 0.02 Hz with 20 ms pulse width, with power of 18 mW/mm^2^. Peak evoked ΔF/F values were measured within a 1 sec time window after stimulation, and average response was calculated. Bicuculline which is a GABAA receptor antagonist was also tested both individually and also in a combination with two glutamatergic blockers. D-AP5 which is an NMDA receptor antagonist, and NBQX which is an AMPA receptor antagonist were used as blockers of glutamatergic synaptic activity. Similarly, 10 pulses at 0.02Hz with 20ms pulse width were delivered and peak values were measured within a 1sec time window post stimulation. The average response of 10 pulses in the presence of bicuculline only and all drugs conditions were recorded. Values were measured from the ROI exactly under the stimulation area.

### Pharmacology

Bicuculline (Tocris) was dissolved in distilled water for final concentration of 10 μM. NBQX was dissolved in dimethyl-sulfoxide (DMSO) to achieve 50 µM final concentration and D-AP5 was dissolved in phosphate buffered saline (PBS) to get final concentration of 100 µM.

### Immunofluorescent Staining

Cultures were fixed in 4% Paraformaldehyde for 2 hours, removed from substrate, washed and permeabilized with 0.3% Triton X-100 (Sigma-Aldrich) in phosphate-buffered saline (PBS) for 2 hours on a shaking platform at RT. Cultures were then blocked in 10% goat serum in PBS for 1 hr. All the primary antibodies were applied to the culture for 48 hours on a shaking platform at 4 °C. Anti-NeuN conjugated to Alexa Fluor 555 (MAB377A5, Millipore) was applied at 1:500 dilution. Anti-Beta-III Tubulin (ab78078, Abcam) was at dilution of 1:1000. Alexa Fluor 488 was used as secondary antibody for Anti-Beta-III Tubulin at dilution of 1:1000. Optical stacks were imaged from the entire thickness of the cultures using Zeiss confocal microscope with 40X or 25X objectives (Zeiss LSM 880, Germany). Optical slices were taken with 1 μm intervals. Images were then processed in ImageJ. For neuronal counting, we slightly modified the existing 3D watershed technique from^57^ to detect the nuclei of pyramidal cells in CA1 and CA3 layers and granule cells in DG neuronal layer.

### Statistics

Statistical tests were used as described in the Results section. Nonparametric tests were used for data that were not normally distributed. A statistically significant difference was defined as *p* < 0.05. At least 3 biological replicates were analyzed for all experiments. The number of biological and technical replicates for different experiments is indicated in the Results section. No specific a priori calculation of sample size was performed. No data or outliers were excluded.

## Data Availability Statement

Figure source data files contain the numerical data used to generate the figures.

## Code Availability Statement

Matlab code and data for the two computational models developed for this study are available on Zenodo via GitHub^50^.

## Author Contributions

SG, ZIF, and YB conceived the work, designed the experiments, and wrote the manuscript. SG performed the organotypic culture experiments, analyzed the data, and devised the spherical spiking model. ZIG performed dissociated culture experiments, and analyzed the data. YB devised the rate-based model, supervised the experiments, result interpretation and spherical spiking model formation. MB and Md.FH maintained organotypic cultures and acquired recordings.

## Acknowledgements

The research reported in this publication was supported by the National Institute of Neurological Disorders and Stroke of the National Institutes of Health under Award Number R21/R33NS088358. This research was also supported in part by National Science Foundation under Award Number NCS ECCS 1835278. The content is solely the responsibility of the authors and does not necessarily represent the official views of the National Institutes of Health or the National Science Foundation.

## Competing Interests

The authors declare no competing interests.

---

**Figure 2 – figure supplement 1.**
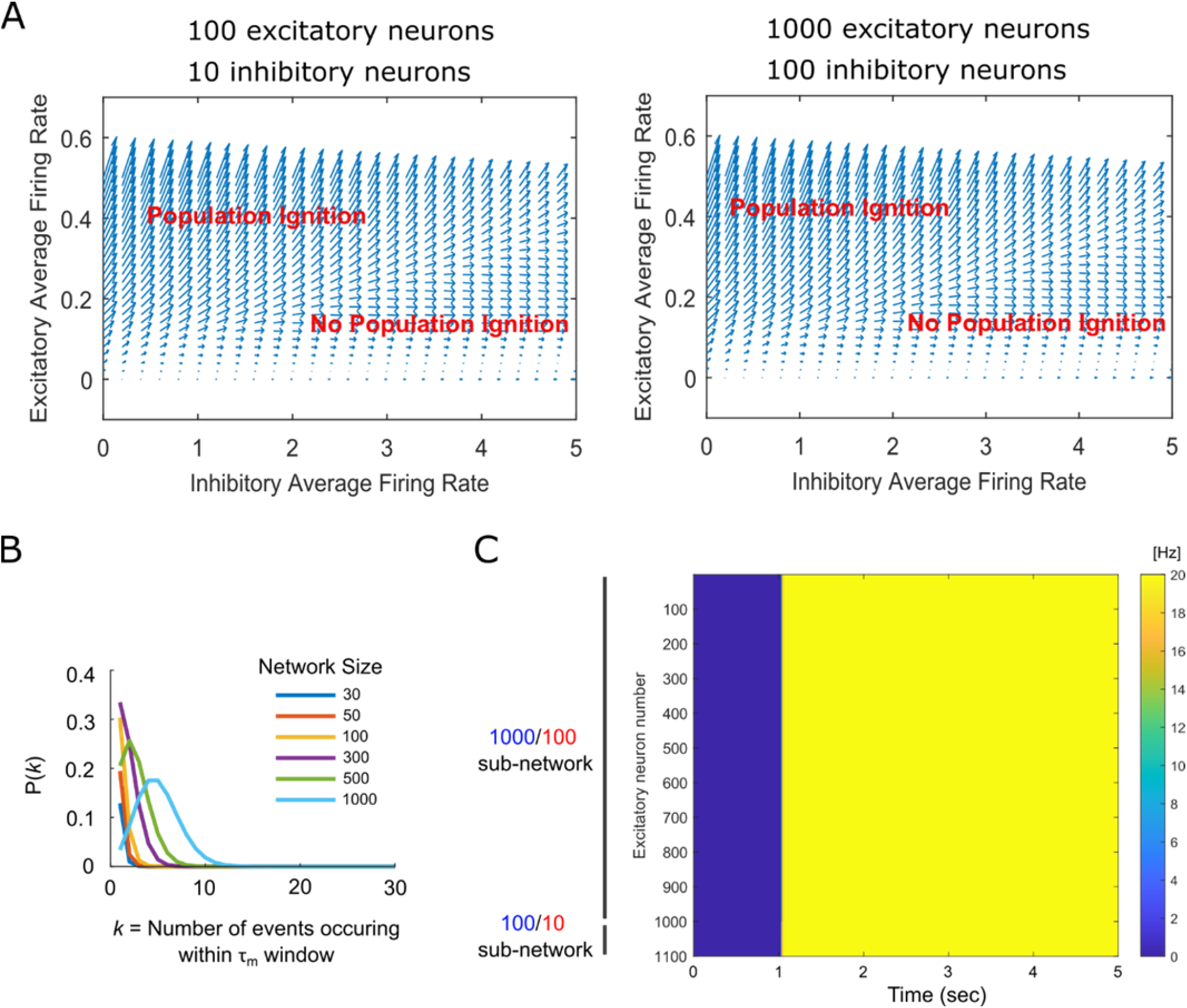
Population ignition in rate-based model. (**A**) Representative phase portraits of 100/10 and 1000/100 networks. Each arrow represents the direction of the trajectory at that point. Points within ‘Population Ignition’ region indicated that a system with that average excitatory and inhibitory rate will achieve population ignition. Conversely, points within ‘No Population Ignition’ region indicate that the system will recover to zero firing rate without population ignition. (**B**) Probability of *k* events of a 0.5 Hz Poisson noise event process occurring within a τ_m_ time window for networks of different size. Probability of population ignition is calculated from this data by determining the probability of *k* > minimum number of neurons for ignition for network of corresponding size. (**C**) Representative raster plot of successful ignition of a large sub-network (excitatory neurons 1-1000) by a small sub-network (excitatory neurons 1000 – 1100).

**Figure 3 – figure supplement 1.**
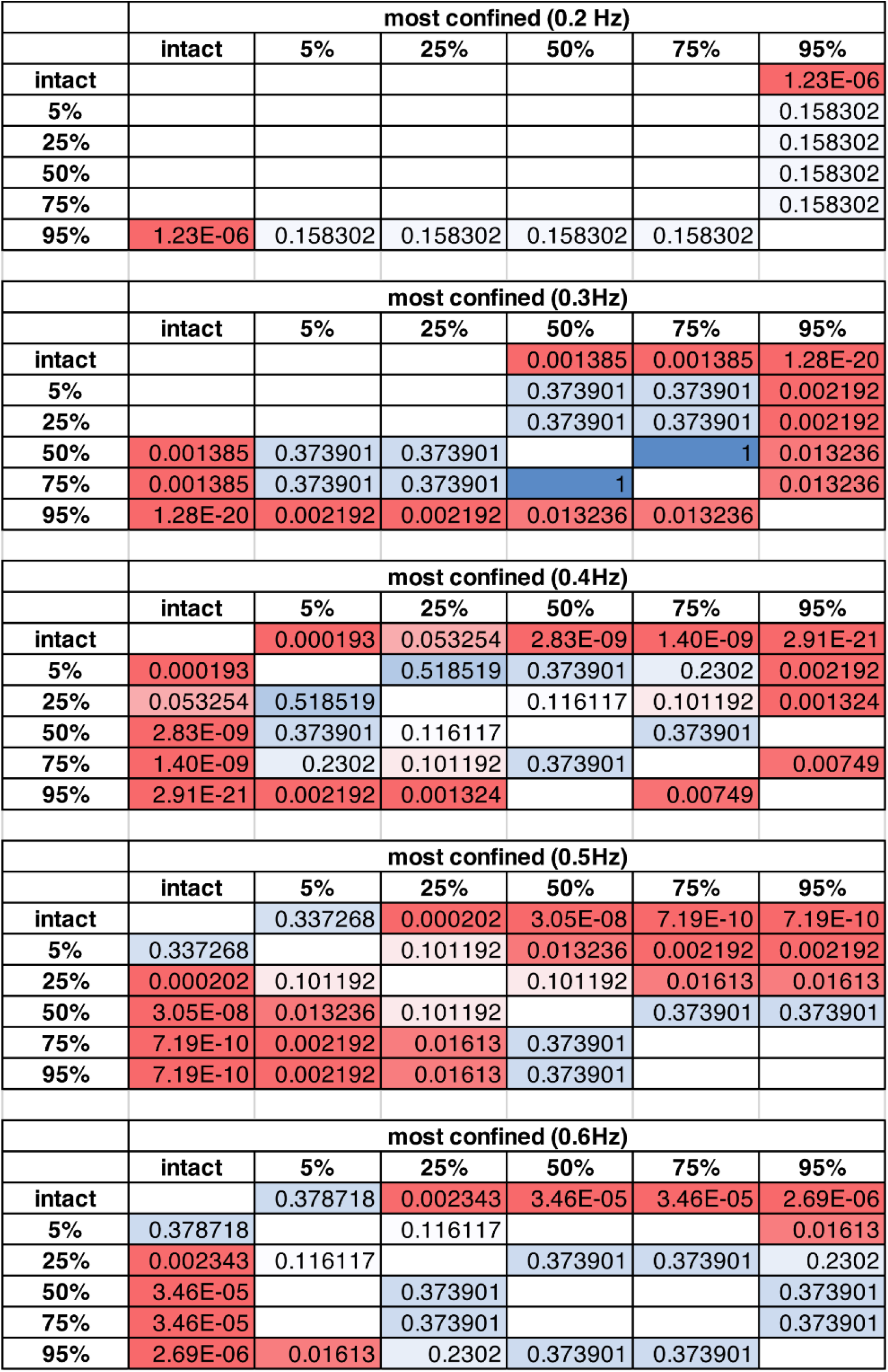
*P*-values (two-sample t-test) for data presented in Figure 3(F): comparison of the number of population bursts in spherical networks with different injury size (%), under the most confined (short sprouting critical distance) constraint.

**Figure 3 – figure supplement 2.**
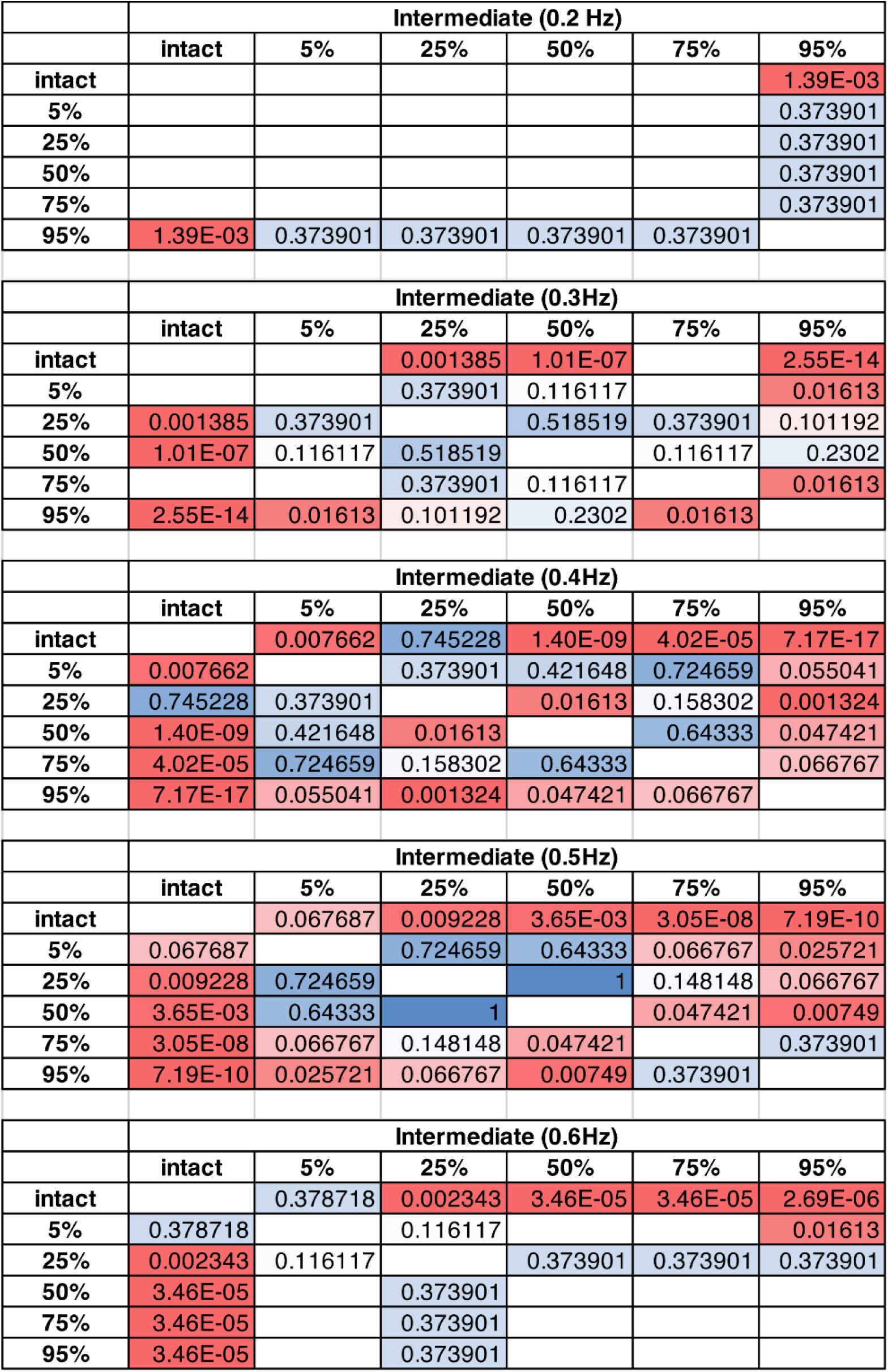
*P*-values (two-sample t-test) for data presented in Figure 3(G): comparison of the number of population bursts in spherical networks with different injury size (%), under intermediate sprouting critical distance constraint.

**Figure 3 – figure supplement 3.**
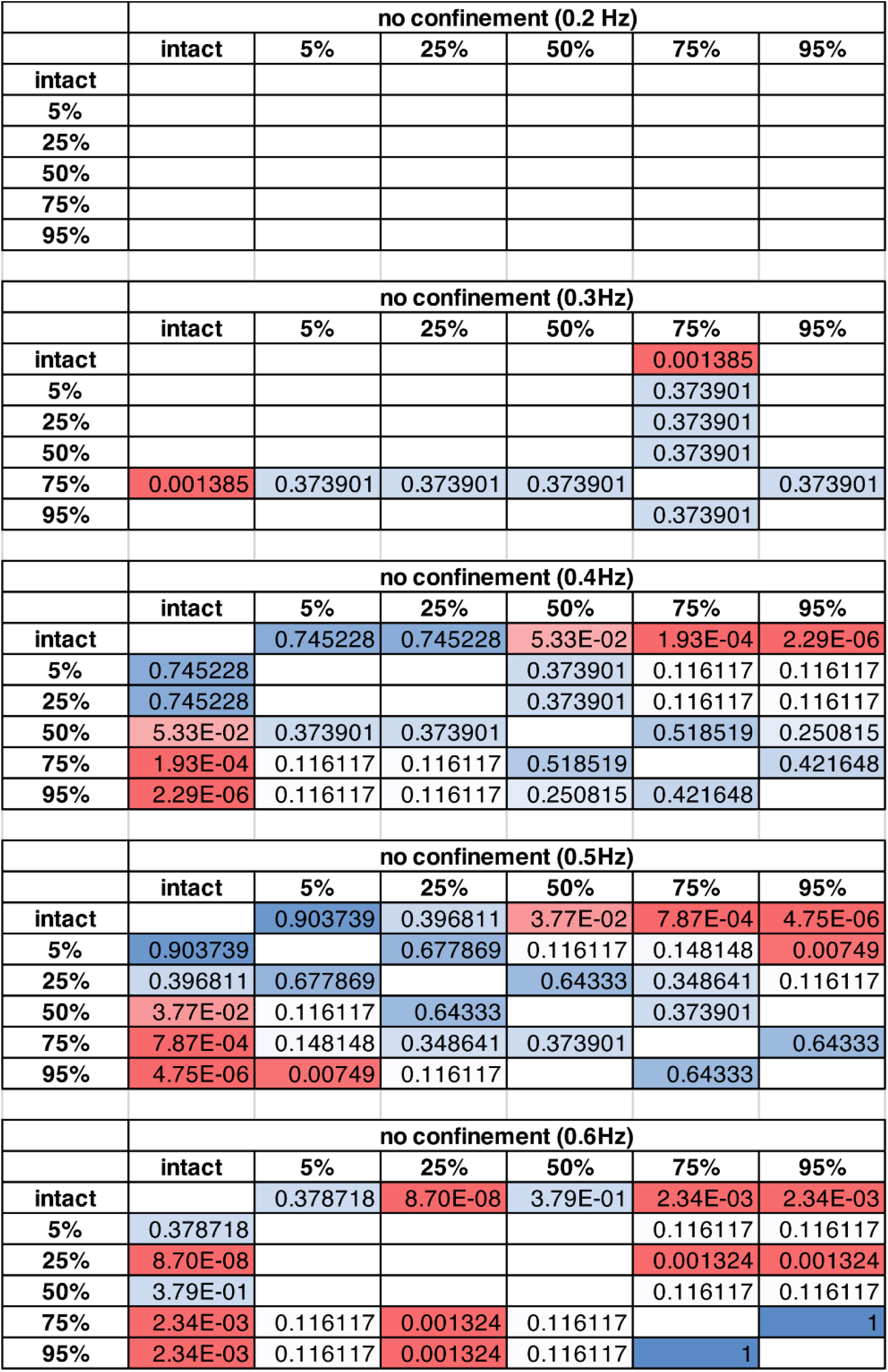
*P*-values (two-sample t-test) for data presented in Figure 3(H): comparison of the number of population bursts in spherical networks with different injury size (%), under no confinement constraint (sprouting critical distance is the same as during network development prior to injury).

**Figure 5 – figure supplement 1.**
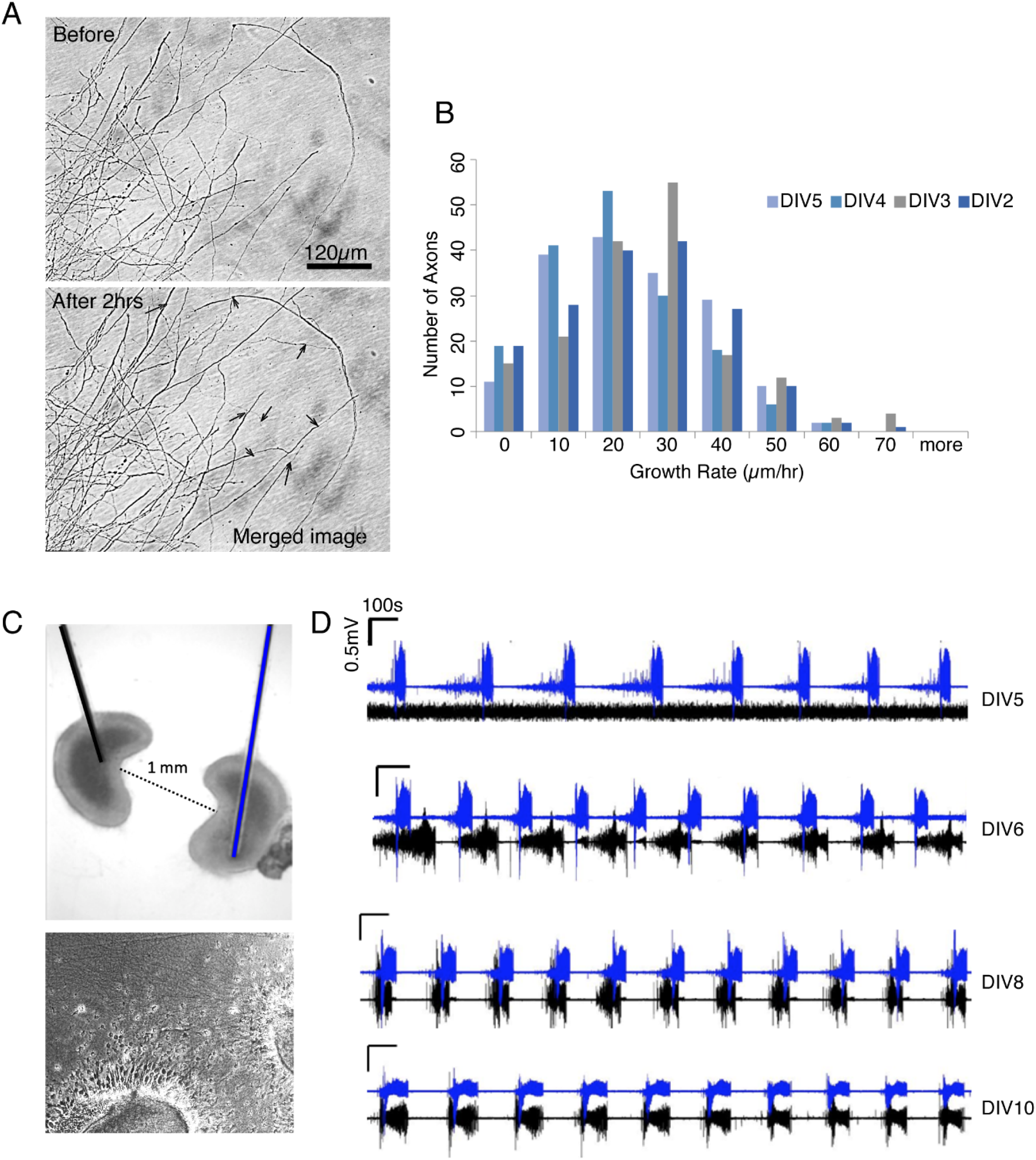
Axon sprouting and functional re-wiring in organotypic cultures. (**A**) Representative phase contrast micrographs of axon sprouting by neurons in organotypic hippocampal cultures. Arrows indicate where axon growth has occurred compared. Scale bar 120 µm. (**B**) Histogram of axonal growth rates. *n* = 3 cultures, 4 days, 170 axons per day. (**C**) Top: bright field of a pair of CA3 slices co-cultured 1 mm apart from each other together on tungsten microwire electrodes (color-coded blue and black). Bottom: phase contrast image of the space between co-cultured slices shows dense axonal sprouting. (**D**) Representative electrical recording of spontaneous activity in co-culture via corresponding electrodes. On DIV 10 ictal-like activity is synchronized.

**Figure 7 – figure supplement 1.**
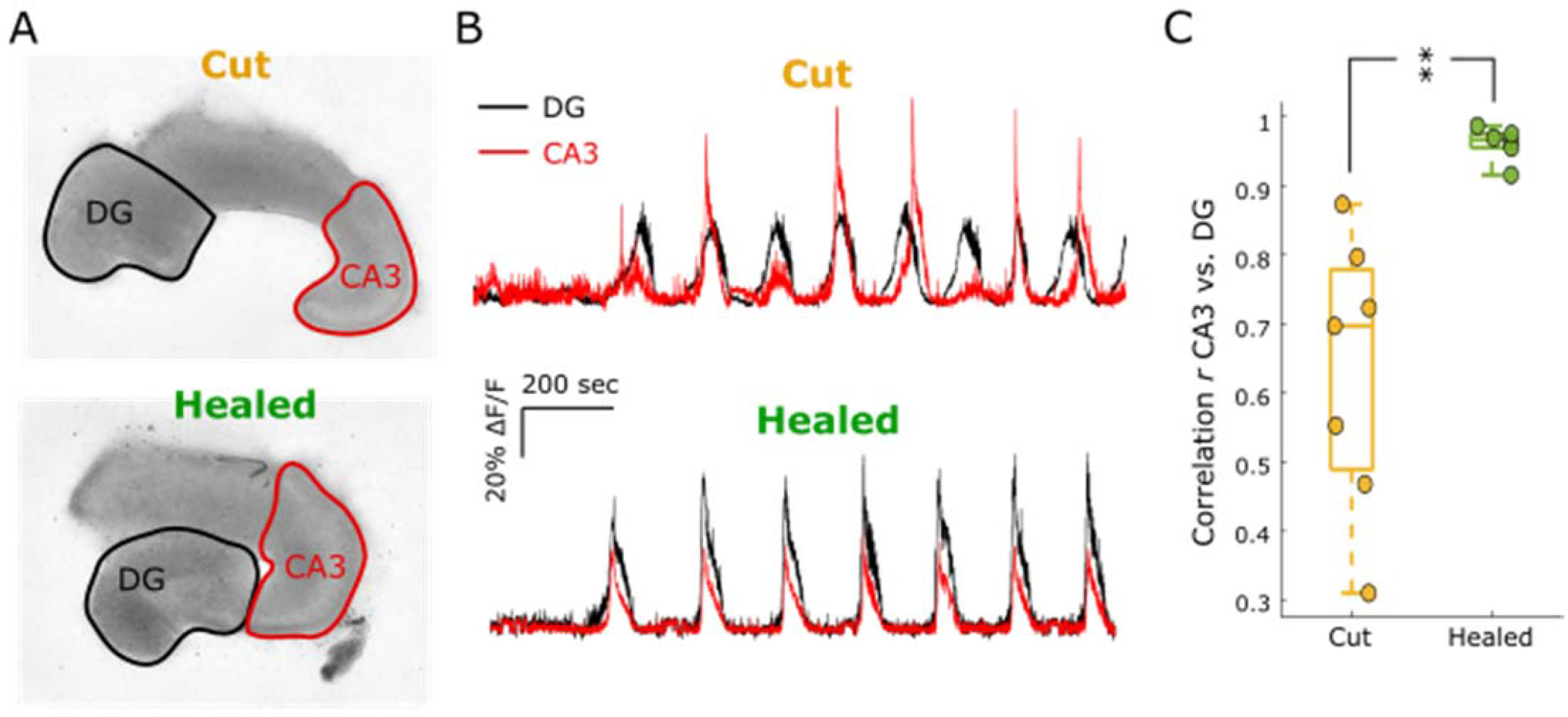
Healing of the cut between CA3 and DG restores activity correlation. (**A**) Phase micrographs of cut and healed cultures, with cut delivered to CA3c. (**B**) Representative traces of spontaneous activity (changes in jRGECO1a fluorescence) in CA3 (red outline in (A)) and DG (black outline in (A)) in cut and healed cultures. (**C**) Plot of the correlation coefficient r of activity in CA3 and DG in cut and healed cultures. ** p < 0.01

**Figure 8 – figure supplement 1.**
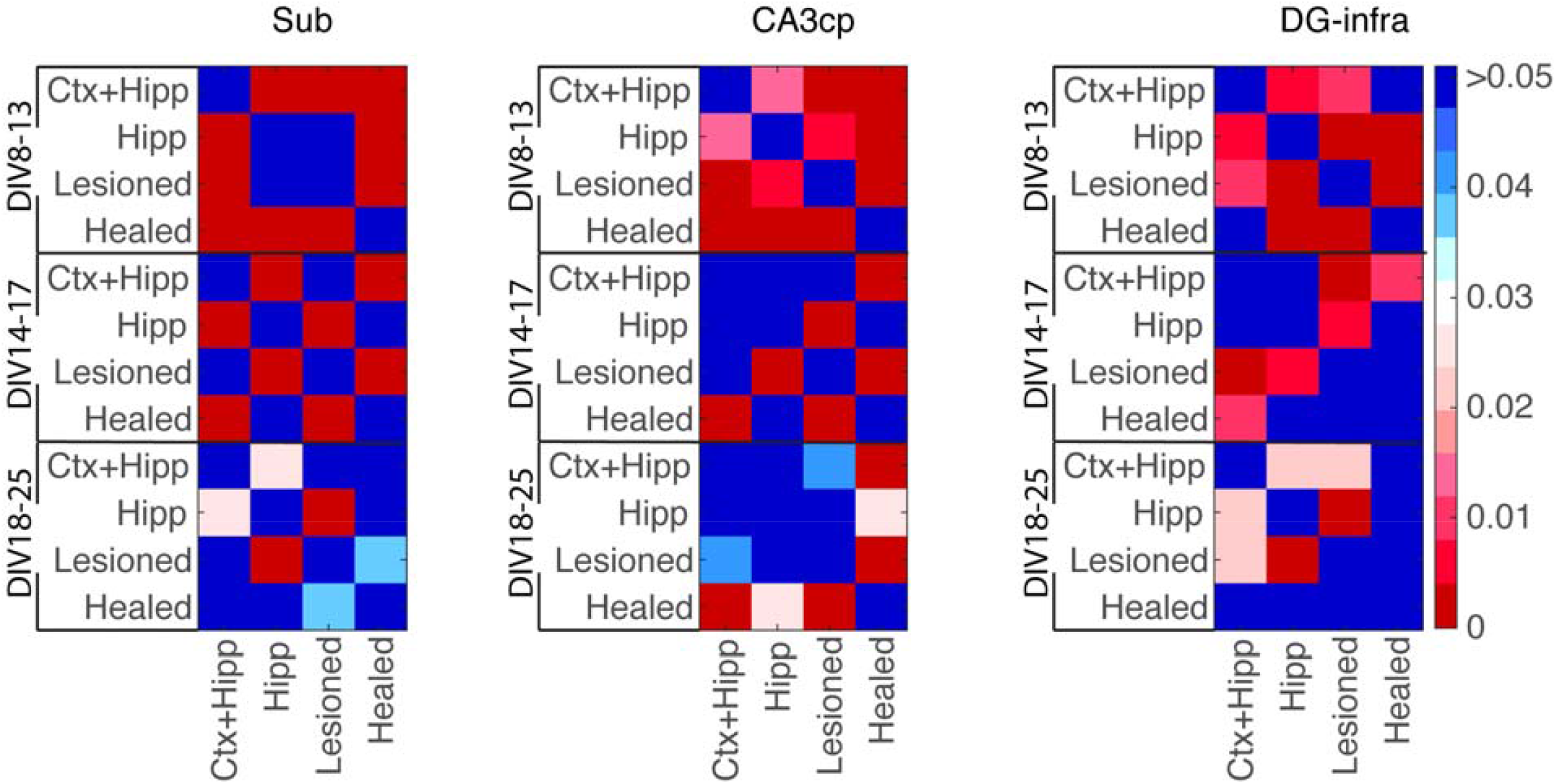
*P* values (population proportion through *z* score) for comparisons of ictal-like activity initiation site; for data presented in Fig. 8 (A-C). DIV8-13: *n*(ctx+hipp) = 35, *n*(hipp) = 51, *n*(lesioned) = 45, *n*(healed) = 12 events. DIV14-17: *n*(ctx+hipp) = 14, *n*(hipp) = 78, *n*(lesioned) = 63, *n*(healed) = 56 events. DIV18-25: *n*(ctx+hipp) = 10, *n*(hipp) = 51, *n*(lesioned) = 51, *n*(healed) = 29 events. Overall, 9 ctx+hipp cultures from 3 animals, 8 hipp cultures from 3 animals, 7 lesioned cultures from 3 animals, and 4 healed cultures from 3 animals were used for this experiment.

## Notes

### Competing Interest Statement

The authors have declared no competing interest.

